# Non-invasive Topographic Sculpting of the Mammalian Auditory Cortex with Low-intensity Focused Ultrasound: A Platform for Personalized Editing of Sensory Representations

**DOI:** 10.64898/2026.11.25.769300

**Authors:** Jacob R. Hehir, YuBing Yolanda Shen, Claire J. Lee, Jonathan A. N. Fisher

## Abstract

**Background:** Topographic maps organize the mammalian cortex, and their maladaptive reorganization underlies conditions like chronic tinnitus and focal dystonia. Low-intensity focused ultrasound (FUS) produces offline neuromodulation, yet its submillimeter spatial distribution within sensory maps remains poorly understood.

**Objective:** To determine how offline FUS-induced amplitude changes are spatially organized within the mouse primary auditory cortex (A1) tonotopic map and depend on sonication intensity.

**Methods:** Widefield Ca2+ imaging was combined with phased-array FUS (750 kHz, <1.5 W/cm^2^ *I_sppa_*, no microbubbles) in Thy1-GCaMP6s × CBA/CaJ mice. Twenty-second sonication was steered to each animal’s tone-evoked centroid at 0.625 or 1.2 W/cm^2^. Offline gain was quantified over 40 min using Euclidean distance from the focus and functional distance (first principal component, PC1, percentile brackets).

**Results:** Modulation followed functional rather than physical geometry: effects appeared isotropic in Euclidean distance but varied spatially across functional brackets, peaking 10 min post-sonication and persisting upwards of 40 min. Both intensities carved a modulatory trough between the 50th and 70th percentiles of A1’s sensory representation (∼1.0–1.2 mm from target). At 0.625 W/cm^2^ *I_sppa_*, this trough became suppressive (gain < 1.0) and shifted relative response timing across cortical fields, an effect absent at 1.2 W/cm^2^.

**Conclusions:** Offline FUS modulation in A1 is organized by functional topography rather than physical distance, demonstrating that low-intensity FUS can serve as a spatially constrained suppressant. A forward-projection model built on this profile provides a quantitative basis for personalized, coordinate-based sonication planning for focal sensory pathologies like tinnitus.

**Highlights:**

- Offline FUS gain in mouse A1 is organized by functional topography, not physical distance
- A shared modulatory trough sits at the 50th to 70th percentile of the tonotopic map
- Intensity sets the sign of offline gain (0.625 suppresses, 1.2 W/cm^2^ *I_sppa_* amplifies)
- Low-intensity sonication can achieve true focal suppression (gain < 1.0) within this trough.
- Relative response timing across auditory cortical regions shifts after low-but not high-intensity sonication
- A forward-projection model supports personalized, coordinate-based sonication planning

## 1. Introduction

The functional organization of the mammalian cortex into topographic maps, in which sensory features are represented by distinct neural populations, is fundamental to perception. Migraineurs with visual auras experience this spatial mapping which manifests as a jagged, colorful border slowly drifting across their visual field. This migrating blind spot (scotoma) is driven by a wave of cortical spreading depolarization traveling across the visual cortex, directly mirroring the wave’s physical path through the brain (Hadjikhani et al., 2001).

Map plasticity, or re-weighting, is equally fundamental to maintaining sensory and motor function (Keck et al., 2008). Acuity and motor control can expand cortical representations through practice, as in musical training (Elbert et al., 1995; Pantev et al., 1998; Pascual-Leone, 2001), Braille reading (Pascual-Leone and Torres, 1993), or acquiring a new spoken language (Menning et al., 2002). The same plasticity also permits maladaptive remapping: phantom limb pain following amputation (Duarte et al., 2020; Lotze et al., 2001), contralesional hemisphere overactivity (Buetefisch, 2015) or remapping (Brown et al., 2007; Dijkhuizen et al., 2001; Sigler et al., 2009) after stroke, altered tactile mapping in focal dystonia (Elbert et al., 1998), and chronic tinnitus following damage to the auditory periphery (Elgoyhen et al., 2015).

Noninvasive neuromodulation offers therapeutic potential for central nervous system conditions that respond poorly to pharmacological treatment. Transcranial magnetic stimulation, direct and alternating current stimulation, and, more recently, temporal interference stimulation (Grossman et al., 2017; Violante et al., 2023) use magnetic or electrical fields to modulate cortical activity. Several of these modalities are in clinical use, but their spatial resolution is coarse relative to primary sensory maps, which are compressed to the millimeter scale: human tonotopy spans roughly 27 to 30 mm across seven octaves in Heschl’s gyrus (Da Costa et al., 2011), and the equivalent gradient in mouse A1 occupies 1 to 2 mm (Tsukano et al., 2016).

Low-intensity FUS restricts modulatory stimuli to volumes several orders of magnitude smaller, and to a substantially finer focal cross-section, than transcranial electrical or magnetic methods (Blackmore et al., 2019). In humans, low-intensity transcranial FUS has been used to modulate cortical and deep subcortical targets while simultaneously capturing the effects via neurochemical and connectivity imaging (Yaakub et al., 2023). It has also been used to engage pain circuits (Riis et al., 2024) and to deliver depth-specific, priming protocols to the motor cortex that leverage metaplasticity (Bao et al., 2024; Ding et al., 2024). The modulatory effect and its dependence on acoustic parameters are not simple functions of membrane biophysics; rather, they reflect cell-type-specific sensitivities (Murphy et al., 2022; Yu et al., 2021). Consequently, altering the intensity and pulse regime can selectively elicit or suppress spiking to up- or down-regulate network activity.

Most FUS research has used the modality for transient activation or acute suppression, but much of its clinical relevance lies in “offline” effects, defined as modulation that outlasts the sonication period itself (Verhagen et al., 2019). Offline effects have been reported to persist for 10 to 20 min after sonication in rodent and primate preparations (Dallapiazza et al., 2018), with more pronounced effects at higher intensities and duty cycles (Kim et al., 2014), and they are the regime in which most human FUS neuromodulation studies operate (Kim et al., 2014; Riis et al., 2024; Yaakub et al., 2023). Restricting measurement to this regime also removes the transient peripheral auditory activation that accompanies active sonication and confounds online readouts (Braun et al., 2020; Guo et al., 2018; Mohammadjavadi et al., 2019; Sato et al., 2018), a consideration of particular weight in auditory cortex.

Toward developing a calibrated intervention, the spatial footprint of the resulting effect must be known, i.e., where, relative to the intended target, gain rises or falls, and by how much. What is currently known about that footprint for offline effects comes largely from macroscale imaging, which integrates millimeters to centimeters and treats tissue volume as its unit. A robust finding is that sonication alters not just the sonicated volume, but distal regions connected both functionally and structurally to the targeted region (Pellow et al., 2024; Verhagen et al., 2019; Yang et al., 2022a, 2022b, 2021). In contrast, off-target effects of transcranial magnetic stimulation (TMS) may be preferentially structurally connected, based on correlation with white matter structures discerned by diffusion MRI (Momi et al., 2021). In both cases the affected territory is set by connectivity rather than by proximity. Connectivity is also what organizes cortex at the millimeter and sub-millimeter scale: horizontal connections within a cortical area, and in sensory areas particularly, are laid out with respect to the axes of the underlying functional map rather than isotropically (Oviedo et al., 2010; Read et al., 2001; Song et al., 2006; Watkins et al., 2014). Whether offline modulation inherits that organization is untested. The acoustic focus may be sub-millimeter in FWHM beam width, but the sub-millimeter spatial pattern of offline modulation around that focus has not been measured, leaving open how a sonication pattern should be designed to achieve a desired fine-scale sculpting of a sensory map.

Auditory cortex is a structurally, functionally, and translationally opportune region in which to explore this question in animal experiments. Tonotopy is preserved across mammals (Bowen et al., 2020; Da Costa et al., 2011; Romero et al., 2020; Tsukano et al., 2016) and can be mapped in small animal models with Ca^2+^ (Romero et al., 2020) or optical intrinsic signal imaging, including marmosets (Zeng et al., 2019), and using fMRI in nonhuman primates (Petkov et al., 2006) and humans (Leaver and Rauschecker, 2016; Moerel et al., 2014).

The system lends itself to quantitative psychophysics behavioral assays given that a position in that map represents an audible tone, and the tone is also what the listener reports: at roughly 4 mm of human cortex per octave, a sub-millimeter shift in representation spans a frequency interval nearly two orders of magnitude larger than the pure-tone difference limen (Micheyl et al., 2012; Wier et al., 1977), and behavioral results can be assessed by standard audiometry protocols. Similarly, tinnitus represents a neuropathology directly mapped to this tonotopic architecture. Clinical pitch-matching of the phantom percept corresponds to focal functional alterations in human neuroimaging (Elgoyhen et al., 2015). This spatial reorganization is corroborated by animal models, where mice with noise-induced high-frequency hearing loss develop an expanded representation of spared lower-frequency tones in A1 (Willott et al., 1993), mirroring the maladaptive plasticity characteristic of human tinnitus.

We sought to establish the spatial reference frame governing offline FUS neuromodulation at the sub-millimeter scale. By integrating *in vivo* widefield Ca²⁺ imaging with functionally targeted low-intensity FUS in mouse primary auditory cortex, we tracked how sonication alters cortical gain and response timing across the broader sensory map. Rather than assuming neuromodulation spreads symmetrically from the acoustic focus, we evaluated the resulting plasticity across two competing coordinate systems: physical distance from the acoustic target versus functional distance within the subject’s own tonotopic map. We used this approach to define the spatial rules of FUS-induced plasticity and provides the empirical foundation necessary to build predictive models for personalized, functionally guided sonication.

## 2. Materials and methods

### 2.1 Animals and surgical procedures

Adult male Thy1-GCaMP6s (GP4.3) x CBA/CaJ mice 4–6 months of age were used in all experiments. The cross The GP4.3 line has a hearing-loss phenotype that can be rescued by breeding with preserved-hearing lines such as CBA/CaJ (Bowen et al., 2020). Hearing was confirmed to be preserved in all mice used, through ABR testing (Fig. S1). Mice were anesthetized with an induction dose of 5% isoflurane (vaporized in medical grade compressed oxygen) at 2 L/min and maintained with 1.5 - 2% isoflurane at 2 L/min, and were given a bolus of buprenorphine (0.5 mg/kg, subcutaneously) and dexamethasone (0.2 mg/kg, subcutaneously). Temperature was maintained at 37°C with a closed-loop heating pad and rectal probe (40-90-8D, FHC, Inc., Bowdoin, ME). Petrolatum ophthalmic ointment (Puralube, Fera Pharmaceuticals, Locust Valley, NY) was used to protect the eyes. The skull was scored with a scalpel to improve head-mount adhesion; a custom aluminum bar was affixed lateral to the sagittal suture with epoxy (Loctite 454) and dental acrylic (C&B Metabond, Parkell Inc., Edgewood, NY). Skin and temporalis muscle overlying the right temporal skull were removed, and a ∼4 x 3 mm craniotomy was made with a dental drill (EXL-M40/LHP12, Osada Inc., Los Angeles, CA), bounded by the zygomatic arch (ventral), temporal ridge (dorsal), lambdoid suture (caudal), and squamosal suture (rostral).

Cranial windows were sealed with a 2.5 × 2.5 mm square piece of poly(4-methyl-1-pentene) (PMP) (Goodfellow Cambridge Ltd, Huntingdon, England) stored in 70% ethanol. Optically clear silicone elastomer (Kwik-Sil, World Precision Instruments, FL) was applied to the exposed cortex via a mixing-tip applicator (World Precision Instruments, Sarasota, FL) until it impressed but did not occlude the cortical surface. Loctite 454 (HenKel, Wisconsin) sealed the window edges, solidified with Loctite Accelerator (SF 712), and dental cement (C&B Metabond, Parkell, New York) covered all exposed skull and window periphery. Mice were allowed to recover for 5 days before imaging. Animals were euthanized directly following imaging sessions via cervical dislocation.

### 2.2 Widefield Ca^2+^ imaging

We used a custom opto-acoustic widefield imaging system that integrated a multi-zoom fluorescence macroscope (Nikon AZ100) together with ring transducer mounted on the objective. The use of a long working-distance objective (5X, AZ-Plan Fluor, numerical aperture: 0.5; working distance: 15 mm) enabled the acoustic and optical foci to co-localize. Illumination was provided by an external halogen lamp (Lambda LS, Sutter Instruments, Novato, CA) filtered to 470 ± 20 nm and delivered via liquid light guide, gated by a shutter (Uniblitz VMM-D1, Vincent Associates, Rochester, NY). Images were acquired with a CMOS camera operating at 16-bit depth (Zyla, Andor Technology Ltd., Belfast, UK) at a sampling rate of 8.57 Hz. All hardware synchronization and triggering used custom LabVIEW programs (National Instruments, Austin, TX).

### 2.3 Optical coherence tomography angiography (OCT-A)

Vascular integrity was assessed with a custom spectral-domain OCT system, described previously (Shen et al., 2025), incorporating a superluminescent diode source (Superlum, cBLMD-T-850-HP, λc=850 nm, Δλ=160 nm) and a spectrometer (Cobra-S 800, Wasatch Photonics) with a line-scan camera (Teledyne e2v) at 80 kHz. A 5X, long-working-distance objective (M Plan Apo NIR, Mitutoyo) achieved axial and lateral resolution of ∼2.5 μm and ∼3.5 μm in tissue, respectively. Custom C++/CUDA software controlled acquisition and performed near-real-time GPU processing (GeForce RTX 3090, NVIDIA), including CUDA-accelerated DFT registration and split-spectrum amplitude-decorrelation angiography to isolate flowing-blood signal from static reflectance. Each volumetric snapshot comprised 600 × 600 cross-sectional planes; repeated B-scans were acquired per location. Total acquisition and processing time per angiographic volume was 30 s. *En face* projections were generated from flattened OCT-A images in MATLAB (MathWorks, Natick, MA).

### 2.4 Immunohistochemical procedures

Immunohistochemistry followed previously described methods (Shen et al., 2025) with minor modifications. Following dissection, tissue was post-fixed in 4% paraformaldehyde (24 h, 4°C), rinsed in PBS (1 h), and equilibrated in 30% sucrose. Tissue was embedded in OCT compound (Tissue-Tek, Sakura Finetechnical Co., Tokyo, Japan) and cryosectioned at 18 μm. Sections were blocked (10% normal goat serum [NGS], 0.4% Triton X-100 in PBS, 1 h, room temperature) and stained overnight at 4°C with CD31 rat monoclonal antibody (PECAM-1; 1:500; Abcam 56299) in 2% NGS/0.3% Triton X-100, followed by Alexa-488-conjugated goat anti-rat secondary (1:500, Abcam 150165, 1 h). Sections were mounted, dehydrated, and coverslipped with Krystalon non-fluorescent medium (64969-95, EMD, Darmstadt, Germany). Fluorescence microscopy used a Keyence BZ-X810 All-in-One system (Osaka, Japan); Evans blue was imaged at 560 ± 28 nm (excitation) / 645 ± 38 nm (emission), CD31 at 470 ± 40 nm / 525 ± 50 nm. Blood-brain-barrier (BBB) integrity was assessed via extravasation of Evans blue dye (0.1 mL of 1% solution, i.p., 30 min before FUS), using the same sonication parameters as for CD31 distribution analysis. Post-treatment, brains were harvested, fixed, and processed as above.

### 2.5 FUS sonication protocol

A custom 16-channel phased-array transducer (H-405, Sonic Concepts, Inc., Bothell, WA) steered the acoustic focus; target coordinates were converted to per-element phase adjustments with custom MATLAB code. Beam-steering accuracy was verified *in situ* using a thermochromic film fragment (35 to 40°C sensitivity) at 100% duty cycle, imaged with a color camera (DS-Fi3, Nikon Corp., Tokyo, Japan) (Fig. S2). Analog waveforms were generated by a high-power driver (TPO-205-07, Sonic Concepts, Inc.). For neuromodulation, pulsed 750-kHz FUS was applied at 5-ms burst length and 100-Hz pulse repetition rate (50% duty cycle) for a total treatment duration of 20 s, at a nominal *I*_sppa_ of either 0.625 or 1.2 W/cm^2^. High-intensity marker doses (>200 W/cm^2^) confirmed target locations via post-hoc histological assessment of vascular integrity and BBB permeabilization. Intensity and beam profile were calibrated with a 0.5-mm capsule hydrophone (HGL-0200, +3-dB range 0.25 to 40 MHz, 200-μm electrode aperture; Onda Corporation, Sunnyvale, CA) in degassed water. Acoustic energy was coupled to the cranial window through a fluid-filled membrane with ultrasonic transmission gel (Aquasonic 100, Parker Labs, Fairfield, NJ), centrifuged (5,000 rpm, 10 min) to remove bubbles; membrane fluid was degassed for 1 h before use.

### 2.6 Auditory stimulation

Auditory stimuli were delivered through an MF1 multifield speaker (Tucker Davis Technologies) in closed-field configuration via a 10-cm Tygon tube (1/8 inch outer diameter, 1/16 inch inner diameter) inserted into the ear canal contralateral to the craniotomy, powered by an RZ6 processing unit (Tucker Davis Technologies). The speaker was calibrated before every experiment with SigCal software (200 kHz sampling; 5 to 33 kHz calibrated range) to account for ambient temperature and tube-position changes. 50-ms tone pips were tapered with a cos^2^ function, and delivered at 70 dB SPL. Auditory stimuli were delayed relative to the illumination shutter to mitigate cross-modal visual artifacts.

### 2.7 Auditory brainstem response (ABR) testing

All animals were pre-screened for auditory integrity via ABR using the tone-delivery hardware described above. Stimuli consisted of tone pip waveforms at intensities from 20 dB SPL to 80 dB SPL, in 10 dB SPL steps, scanned in 3 dB SPL increments from 5 dB SPL to 33 dB SPL. Mice were anesthetized as in Section 2.1. Three 12-mm, 27-gauge needle electrodes were placed subcutaneously with reference at the midline above the brainstem, active lead either just medial or temporo-lateral to the ipsilateral ear, and ground lead on the proximal hindlimb. Signals were preamplified (Medusa, Tucker Davis Technologies, Alachua, FL) and processed via the RZ6 with BioSig software (Tucker Davis Technologies, Alachua, FL).

### 2.8 Image processing and trace extraction

Raw image stacks were spatially smoothed (10-pixel 2D Gaussian filter). For each pixel, baseline fluorescence (F_0_) was the time-averaged mean of a 270-ms pre-stimulus window, and fractional fluorescence change (ΔF/F_0_) was computed pixel-wise. Principal component analysis (PCA) was used to distill maps of tonotopy and to systematically define spatial regions of interest (ROIs) for sonication targeting, trial averaging, and quantification of offline gain.

ΔF/F_0_ image stacks were spatially down-sampled and linearized (pixels as variables, frames as observations); following min-max normalization, PCA was applied to the pooled dataset of all trials (pre-FUS, post-FUS and sham) for each animal to extract shared spatial modes. An elbow analysis indicated that >80% of variance was explained by the first three principal components (PCs), which were reconstructed into 2D spatial maps (Figs. 1, 2, 4). These PC definitions were used to quantify both response magnitude and spatial distribution.

**Figure 1.**
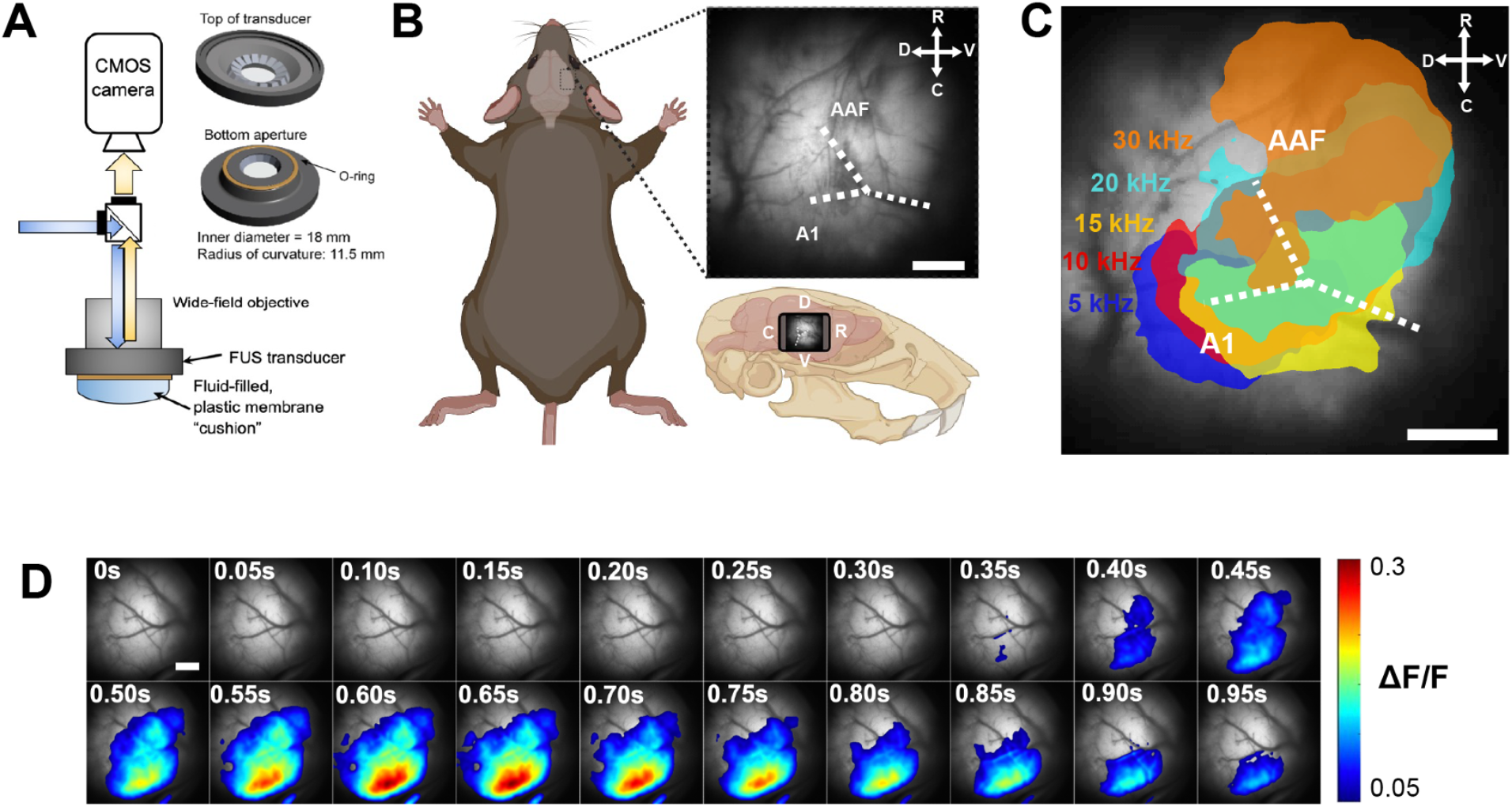
Widefield Ca^2+^ imaging of mouse auditory cortex identifies tone-responsive regions for targeted FUS. (A) Imaging and sonication apparatus (insets: top and bottom of the ring transducer). (B) Location and orientation of the cranial window (bottom) and widefield view of auditory cortex, with primary auditory cortex (A1) and the anterior auditory field (AAF) indicated; the dotted line follows the rhinal vein. (C) Overlay of ΔF/F_0_ responses to 5, 10, 15, 20 and 30 kHz tones in the same field of view; labels as in B. (D) Image sequence of a single-trial response to a 10 kHz tone (onset at 0 s), in 50-ms frames.

**Figure 2.**
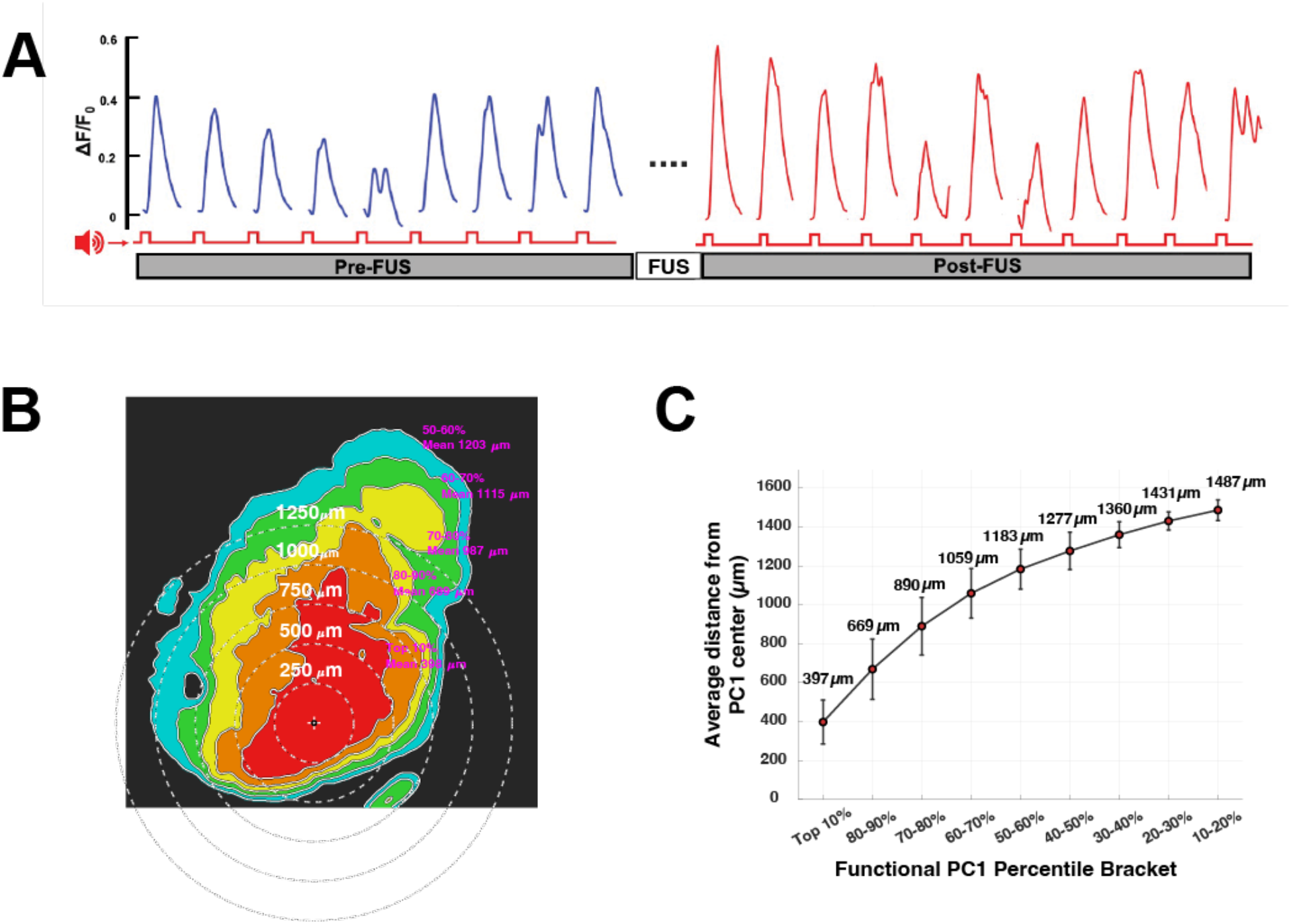
Functional and Euclidean coordinate systems for assessing FUS neuromodulation. (A) Tone-evoked ΔF/F_0_ responses from a representative animal before and after sonication (tone pips, red markers; FUS delivered between the pre- and post-sonication blocks). (B) Pre-sonication PC1 map of the same animal (10 kHz tone-pip stimulus), with functional-percentile brackets (filled contours) and concentric 250-μm Euclidean rings centered on the sonication target. (C) Mean distance (± SD across animals) of each functional-percentile bracket from the PC1 center of mass, which is the sonication target.

One-dimensional temporal traces were extracted by averaging ΔF/F_0_ across spatial masks corresponding to percentile brackets of the normalized PC1 spatial map, generated per experiment. Traces were detrended with a 10-second moving-average filter and baseline-corrected by a secondary mean subtraction. Post-sonication trials were binned into 10-minute windows to assess offline effects while minimizing dropped time points; trials within a block were averaged, and the peak amplitude of the resulting trial-averaged trace was used to calculate gain.

To quantify shifts in the spatial profile of tone-evoked responses following FUS, ΔF/F_0_ image stacks were projected into the PCA subspace and PC1 to PC3 scores were averaged over the evoked-response window.

### 2.9 Data inclusion

A single set of trace-inclusion criteria was applied to all experiments: pre-stimulus stability, defined as a standard deviation within the 270-ms pre-stimulus window with standard deviation ≤ 0.5, and a post-stimulus response, defined as z-score ≥ 5 within a window of 2.42 s following auditory tone presentation, relative to the baseline standard deviation. PCA was performed retrospectively, so a further exclusion criterion was applied post hoc: sonication location, from calibrated steering coordinates, more than 100 μm from the center of mass of the top-10th-percentile PC1 intensity. Of the animals used in this study, one failed to meet this criterion and was excluded from grand averages.

### 2.10 Statistical analysis

Gain was analyzed in two coordinate systems applied to the same measurements (Fig. 1C, 2B). In functional coordinates, cortical location was defined by descending percentile brackets of the normalized PC1 map (top 10% through 10 to 20%); in Euclidean coordinates, by concentric 250-μm annuli centered on the sonication target. For each coordinate system, intensity and 10-min post-sonication window, differences in gain across locations were tested with a Friedman test, with animals as blocks and location as the within-animal factor. Differences between pairs of locations were evaluated with paired t-tests (uncorrected), gain at each location was compared with 1.0 by one-sample t-tests, and gain at each location was compared between intensities by two-sample t-tests (Fig. 2D, E). The spatial distribution of gain across functional brackets was compared between the two intensities with two-way ANOVA tests. Significance was set at P < 0.05.

### 2.11 Spatial gain forecasting via anisotropic tensor warping

To model the continuous spatial distribution of FUS-induced gain across the cortical surface from sparse empirical data, we used a custom spatial predictive pipeline developed in MATLAB (Fig. 5; Fig. S5). Rather than assuming isotropic propagation, which forces radially symmetric footprints, the spatial domain was deformed to respect the irregular architecture of each animal’s baseline cortical response map (PC1 of tone-evoked baseline responses prior to FUS).

The FUS target was identified as the intensity-weighted center of mass (COM) of the top-10th-percentile intensity footprint of the normalized PC1 map. For every pixel, a raw Euclidean distance to this COM was computed, then warped using a directional tensor field built from the local spatial gradient of the normalized PC1 map (Gaussian-smoothed, 5-pixel kernel) and the angular alignment between the propagation path and the local tissue gradient, with anisotropy weight α = 2:

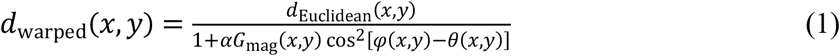

where *d*_Euclidean_ is the Euclidean distance from pixel (*x*, *y*) to the COM, *G*_mag_ is the magnitude of the local PC1 gradient, φ is the orientation of the path from the COM to (*x*, *y*), θ is the orientation of the local PC1 gradient, and α is the anisotropy weight.

Sparse empirical gain measurements sampled at discrete radial intervals were fit with a continuous, single-troughed analytical function characterizing the modulatory trough without piecewise artifacts:

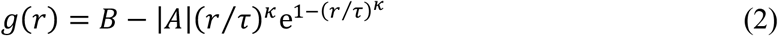

where *B* is the baseline gain, *A* is the peak dip amplitude, τ is the spatial constant defining the trough minimum distance, and κ is a shape and skewness parameter. At κ = 1 this reduces to a standard alpha function; κ ≠ 1 generalizes it to an asymmetric profile, and we refer to it throughout as a generalized (skewed) alpha function. Optimization used a bounded Nelder-Mead simplex search (fminsearch) minimizing a penalized sum-of-squares loss. The optimized 1D profile *g*(r) was evaluated across the 2D tensor-warped distance matrix (converted using a calibrated pixel pitch of 3.24 μm/pixel), and smoothed with a 50-μm Gaussian kernel.

### 2.12 Point-constrained forward-projection optimization for treatment planning

To identify sonication coordinates predicted to produce a specified gain at a chosen cortical location, the forecasting model of Section 2.11 was run forward from candidate foci (Fig. 5 & S5). The warped distance field was generalized to an arbitrary candidate focus (*c*_x_, *c*_y_). Candidate foci were swept across a downsampled grid spanning the field of view (stride, 6 pixels). For each candidate, the gain field *g*(*d*_warped_(*x*, *y*; *c*_x_, *c*_y_)) was computed, and the gain delivered to the chosen location (*G*_projected_) was sampled by linear interpolation. The absolute error relative to the specified gain (*G*_target_) was

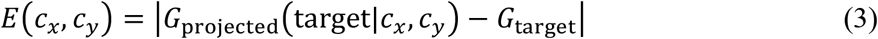

and the error surface was interpolated to full image resolution. Coordinates whose error fell within an error tolerance of

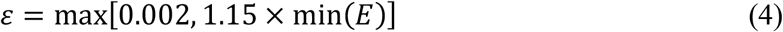

formed the solution arc: a set of approximately equivalent sonication coordinates, each predicted to deliver the specified gain at the chosen location. The optimization used an anisotropy weight of α = 0.8 rather than the value of 2 used for forecasting, because α = 2 produced discontinuous error surfaces that prevented convergence of the search. A single-location optimization was used in place of a global inverse solution for an arbitrary gain landscape, because the radius of the modulatory trough (about 1.15 to 1.2 mm; Section 3.5) occupies a large fraction of the imaging field, which would force candidate foci to the edge of the field. Custom code for this and other analyses is available on GitHub (see Code Availability).

### 2.13 Analysis of principal component response latencies

Response latency was defined as the time from auditory stimulus (the 5-30 kHz) tone pip onset to the peak of the trial-averaged ΔF/F_0_ trace, extracted separately within ROIs given by the top 10th percentile of each of the first three principal components (the PC1, PC2 and PC3 regions) (Section 2.8). Latencies were normalized within animal to that animal’s own pre-FUS value, so that all reported values are fold-changes with pre-FUS defined as 1.0, and were binned into the same 10-minute post-FUS windows used for the gain analysis. Two families of paired comparisons were evaluated: comparisons among the three regions (PC1 vs. PC2, PC2 vs. PC3, PC1 vs. PC3) at each time window and intensity, and longitudinal comparisons of each region against its own pre-FUS baseline (Fig. 4). Holm-Bonferroni correction was applied within each family separately.

### 2.14 Structural modeling of auditory pathways intersected by the acoustic beam

Because the focal region of the transducer is elongated along its propagation axis and the mouse auditory cortex is approximately 1 mm thick, sonication targeted at the surface of A1 is likely to deposit acoustic energy in underlying white matter and in fibers entering or leaving the cortex. To gauge potential off-target effects on the auditory pathway due to interception of axonal tracts within a volume defined by a putative neuromodulatory threshold, we intersected the measured acoustic field with anterograde tract tracing data from the Allen Mouse Brain Connectivity Atlas (Kuan et al., 2015; Oh et al., 2014), registered to the Allen Common Coordinate Framework (CCFv3). Modelling was restricted to the right hemisphere to match the right-sided sonication used in all *in vivo* experiments. Two right-hemisphere anterograde injection experiments served as reference pathways: an injection into the right medial geniculate body (MGB; experiment 180520257, drawn from the Connectivity Atlas), used to represent ascending thalamocortical afferents, and an injection into the right primary auditory area (A1; experiment 120491896), used to represent cortico-cortical and intrinsic efferents. The Allen streamlines are computational reconstructions of bulk anterograde viral fluorescence from a single injection per structure. They do not correspond to individual biological axons and are not weighted by tracer fluorescence. We therefore treat each streamline as a sample of pathway geometry, and report results as percentages of all streamlines in each experiment, since the two experiments differ in streamline number (789 for A1, 213 for MGB), rather than as numbers of axons.

We used the official Allen skeletons (neuroglancer mesoscale skeleton repository) as unbroken polylines instead of a custom tracking routine. Preserving each polyline from injection site to terminal point retains its full length and direction, which allows the terminal structure to be named and avoids the truncation of long or curving bundles that stepwise tracking introduces. Vertices were converted from nanometers to CCFv3 micrometers. Each connected component was counted as one streamline; neither experiment contained branched components, so counts are component counts. Allen stores the lower vertex index at the distal end, so polylines were reversed to run from injection to terminal.

For beam registration, the measured two-dimensional hydrophone profile was revolved about its propagation axis, aligned with the inward cortical surface normal at right A1, and translated so that the acoustic peak coincided with the A1 target, mirroring our experimental targeting. Each polyline was resampled to 25 µm spacing (native spacing ∼103 µm) and the beam field was sampled at every resampled point. A streamline was classed as “intercepted” at a given arbitrary threshold in *I_sppa_* if its maximum sampled intensity was at least that fraction of peak. Among intercepted streamlines, those whose terminal point fell below the threshold were classed as passing through the beam (the beam crosses the fiber in passage), and those whose terminal remained inside the isosurface as ending within it.

Because any single intensity cutoff is physiologically arbitrary, given the cellular, vascular, and acoustic factors that govern neuromodulatory efficacy, we used 50% contour as a case example (Fig. 6) and repeated the analysis for contours from 20% to 100% of peak (Table S6), chiefly to assess a general trend of the relationship between threshold intensity and off-target span and diversity. For streamlines that passed through the thresholded core, the terminal structure was assigned from the 100 µm CCFv3 annotation at the final point and grouped as A1, other auditory cortex regions, temporal association area (TEa), visual cortex, thalamus/MGB (posterior intralaminar thalamic nucleus (PIL), suprageniculate nucleus (SGN), peripeduncular nucleus (PP), posterior triangular thalamic nucleus (PoT)), or other. Terminals within ≤ 5700 µm of the mediolateral axis were classed as contralateral. Custom MATLAB code implementing this pipeline accepts any user-supplied beam profile, stereotactic target, and CCFv3 anterograde experiment (see Code Availability).

## 3. Results

### 3.1 Widefield imaging of tonotopy

Widefield Ca^2+^ imaging produced a tonotopic representation of auditory frequencies within the normal hearing range in Thy1-GCaMP6s (GP4.3) **×** CBA/CaJ mice. In a representative mouse (Fig. 1C), individual tone pips from 5 to 30 kHz at 70 dB SPL elicited Ca^2+^ responses (ΔF/F_0_) of approximately 30% in oblong regions within primary auditory cortex, on the hemisphere contralateral to the stimulated ear. Consistent with prior work, spatial patterns evoked by different tones overlapped, with the border and center of mass for each tone’s response distributed along a gradient that increased in frequency ventrally and rostrally, following the dorsal branch of the rhinal vein (Fig. 1B to C). Tone-evoked cortical responses peaked at 681 ± 10 ms (mean ± standard deviation, n = 12) (Fig. 1E), consistent with the slower kinetics of auditory-evoked responses reported previously in Thy1-GCaMP6s mice (Dana et al., 2014; Romero et al., 2020). Trial-to-trial amplitude was variable, likely reflecting in part spontaneous, widespread increases in ΔF/F_0_ migrating rostro-caudally, consistent with slow-wave activity described previously (Ferron et al., 2009).

### 3.2 In situ validation of targeting and safety

The transducer’s beam profile was characterized with a hydrophone in a tank of degassed water (lateral pressure profile FWHM = 1.7 mm), and targeting accuracy was independently confirmed by color imaging of thermochromic film (Fig. S2) and by histology (Fig. S3). High-intensity marker doses (223 W/cm^2^ *I*_sppa_ at 100% duty cycle) were used to create thermally induced tissue fiducials. These produced localized parenchymal damage and blood brain barrier compromise indicated by Evans blue leakage extending to a depth of ∼ 0.4 mm beneath the cortical surface, confirmed by cryosectioning (Fig. S3A).

Targeted sonication with the FUS intensity of 1.2 W/cm^2^ *I*_sppa_ utilized as the upper bound in our *in vivo* experiments produced no evidence of blood-brain barrier disruption, based on the absence of detectable Evans blue extravasation into the neural parenchyma, and the absence of vascular perturbation or astrocytic scarring, which CD31 and GFAP labelling would have suggested (Fig. S3B). *In vivo* OCT-A showed no acute or persistent change in cortical microvasculature at this dose (Fig. S3C).

### 3.3 Gain is organized by functional cortical topography, not physical distance

By extracting the center of mass of dimensionally reduced focal tonal representations, we precisely targeted the acoustic energy to each animal’s unique functional centroid. We measured offline gain, the ratio of post-to pre-sonication response amplitude, across the field of view and expressed the same measurements in two coordinate systems (Fig. 2). Euclidean coordinates divide the cortex into concentric 250-μm annuli centered on the sonication target. Functional coordinates divide it into percentile brackets of each animal’s own PC1 map, the dominant spatial pattern of its tone-evoked response. The functional brackets are oblong rather than circular, but their mean distance from the target increases almost linearly with bracket order (Fig. 2C; Table S1); the two systems thus sample the cortex at comparable scale, yet differ chiefly in shape.

In functional coordinates, sonication carved an annular trough into the map (Fig. 3A-B). At 0.625 W/cm^2^, gain stayed near 1.0 in the brackets closest to the target, declined through the middle of the map and fell below 1.0 between the 50th and 70th percentile brackets, about 1.05 to 1.2 mm from the target, before recovering toward the periphery. At 1.2 W/cm^2^, gain exceeded 1.0 throughout the map, and the same mid-map brackets showed the smallest increase. The spatial distribution of gain differed significantly between the 0.625 and 1.2 W/cm^2^ conditions across the functional brackets.

**Figure 3.**
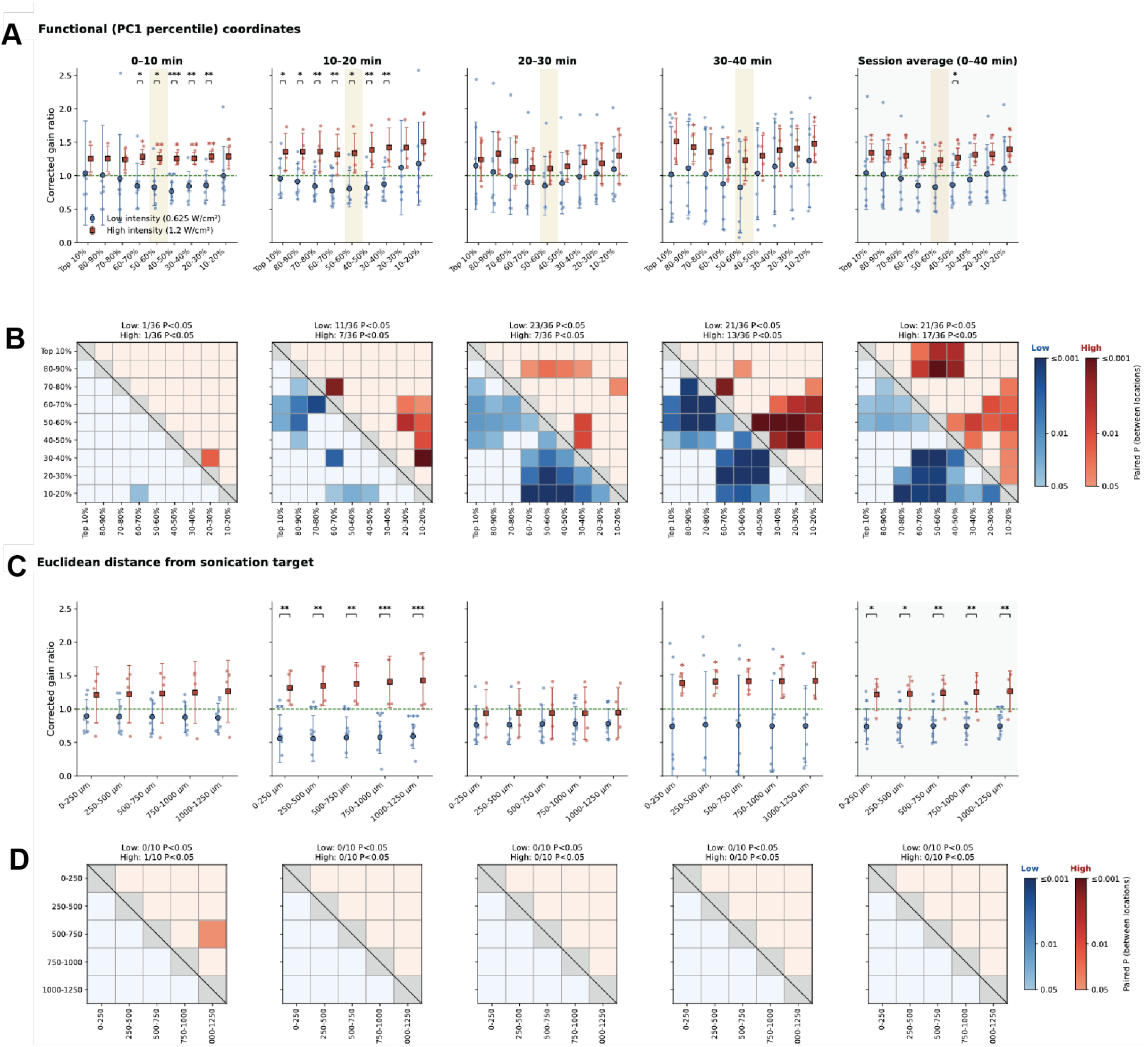
Offline gain is spatially structured in functional but not Euclidean coordinates. (A) Functional coordinates: corrected gain ratio in each PC1-percentile bracket for each 10-min post-sonication window and for the session average (each animal’s mean across the four windows), at 0.625 W/cm^2^ (blue circles) and 1.2 W/cm^2^ (red squares). Small points are individual animals; large markers and error bars are mean ± SD; the dashed line marks a gain of 1.0. Colored asterisks, gain compared with 1.0 (one-sample t-test); black brackets, 0.625 compared with 1.2 W/cm^2^ at that location (two-sample t-test). (B) P values of paired t-tests between all pairs of brackets within each intensity (lower-left triangle, 0.625 W/cm^2^; upper-right triangle, 1.2 W/cm^2^); color saturation encodes P, and pale cells have P ≥ 0.05. The number of pairs with P < 0.05 is given above each matrix; about 1.8 of 36 are expected by chance. (C) Euclidean coordinates, as in Fig. 2B, for 250-μm rings out to 1.25 mm (about 0.5 of 10 pairs expected by chance). (* P < 0.05, ** P < 0.01, *** P < 0.001). 0.625 W/cm^2^, n = 8 animals; 1.2 W/cm^2^, n = 4.

Expressed as Euclidean distance from the target, the same measurements lost this structure. Gain was uniform across all five annuli out to 1.25 mm at both intensities, and only its overall level differed, falling below 1.0 at 0.625 W/cm^2^ and rising above 1.0 at 1.2 W/cm^2^ (Fig. 3C-D). Gain varied across functional brackets in every window after the first at 0.625 W/cm^2^ and with increasing strength over time at 1.2 W/cm^2^, whereas it did not vary across Euclidean annuli in any window at either intensity (Friedman tests: P < 0.0001 at 0.625 W/cm^2^ from 10 min onward; P = 0.051, 0.043 and 0.0032 at 1.2 W/cm^2^ in successive windows; all Euclidean P > 0.3; Fig. S4). Offline modulation therefore appears fundamentally organized by the subject’s own functional architecture, stretching along contours of similar sensory response while ignoring physical proximity.

### 3.4 Temporal evolution of spatial structure

The trough was absent immediately after sonication and developed over the following tens of minutes (Fig. 3A-B). In the first 10 min, gain was flat across the map at both intensities. At 0.625 W/cm^2^, the trough appeared by 10 to 20 min and persisted without decline to the end of the 40-min recording. At 1.2 W/cm^2^, it emerged more gradually and was deepest in the final window. The acoustic field is present only during the 20-s sonication, so the delayed appearance of the trough indicates that its spatial pattern develops through processes that unfold in the cortex after the stimulus has ended, rather than reproducing the acoustic field directly.

Pairwise comparisons locate this development within the map (Fig. 3B,D). In functional coordinates, the number of bracket pairs with differing gain was at chance in the first window and increased thereafter, and the differing pairs consistently set the brackets nearest the target and at the periphery of the map against the mid-percentile brackets that contain the trough. In Euclidean coordinates, no more than one of ten pairs differed in any window (paired t-tests; counts above each matrix in Fig. 3D).

### 3.5 The modulatory trough achieves true sensory suppression at low intensity

To locate the trough precisely and evaluate its absolute depth, we described the gain profile in each window with a single-troughed function of distance from the target (Section 2.11; Fig. S5). Fitted to the windows in which the profile showed spatial structure, the function placed the morphological minimum at 1.12 mm (95% CI 1.10 to 1.14) at 0.625 W/cm^2^ and at 1.15 mm (1.10 to 1.17) at 1.2 W/cm^2^ (Table S2). Both positions fall within the identical functional locus: the 50th to 70th percentile brackets of the sensory map. However, the nature of this trough diverges between the two acoustic regimes. At the high intensity of 1.2 W/cm^2^, the baseline gain across the core of the map was universally elevated (gain = 1.39). The trough at this intensity represents only a relative morphological dip: gain dropped to 1.16 in the mid-map region, constituting a 17% reduction relative to the core, but remaining entirely in a state of net amplification (gain > 1.0). In contrast, sonication at 0.625 W/cm^2^ established a genuine suppressive state. While the core gain remained near unity (1.09), the trough drove the regional gain down to 0.82, a 25% decrease from the local core. Thus, low-intensity FUS can act as a functional depressant tailored to a specific isofrequency domain, providing a mechanism for focal suppression within the tonotopic map.

### 3.6 Low-intensity sonication shifts the relative timing of responses across auditory cortical fields

Tone-evoked activity does not peak simultaneously across auditory cortex. The first three principal components of each animal’s response occupied distinct cortical territories, PC1 centered on A1 and PC2 and PC3 on lateral and anterior auditory regions, each with its own peak latency (Fig. 4). Because FUS alters the coupling between a sonicated region and its distal targets, and not only local excitability (Verhagen et al., 2019; Yang et al., 2022a, 2021), we used these regions to ask whether sonication changes the order in which activity peaks across fields. After sonication at 0.625 W/cm^2^, the peak latencies of the PC1 and PC2 regions diverged within the first 10 min, and those of the PC2 and PC3 regions diverged at 20 to 30 min (Fig. 4; paired comparisons, Holm-corrected P = 0.041 and 0.050; Table S3). Sonication at 1.2 W/cm^2^ produced no such divergence, and at neither intensity did the latency of any region change relative to its own pre-sonication value. Consequently, low-intensity sonication acts not as a global temporal retardant, but as a selective modulator: it alters the relative temporal latency between specific cortical fields while leaving the macroscopic timing of the sensory response intact.

**Figure 4.**
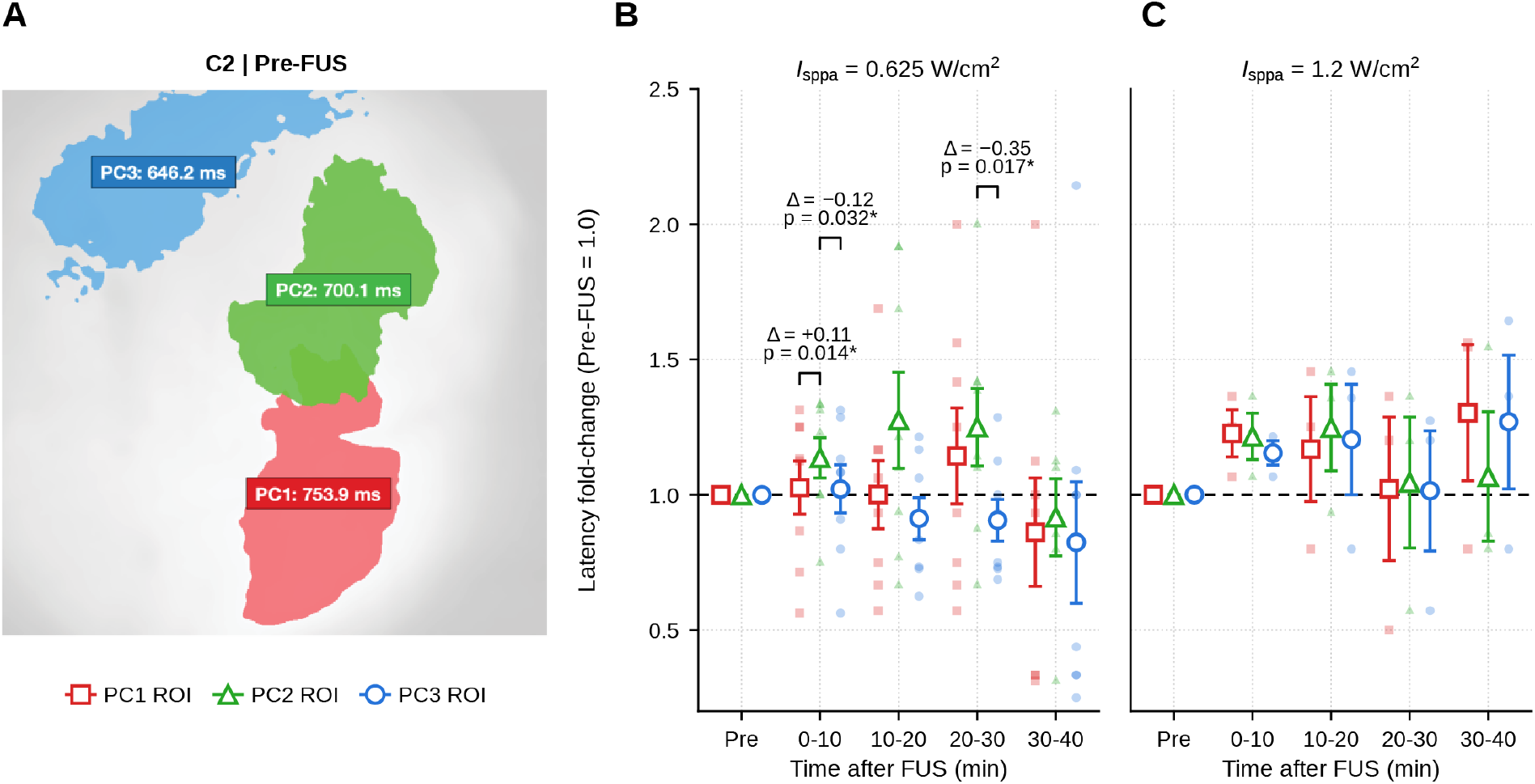
Sonication at 0.625 W/cm^2^ shifts the relative timing of tone-evoked responses across auditory cortical fields. (A) Regions given by the top 10th percentile of PC1 (red), PC2 (green) and PC3 (blue) in a representative animal before sonication, labeled with the time from tone onset to the peak of each region’s trial-averaged response. (B, C) Peak latency of each region relative to its own pre-sonication value (pre-FUS = 1.0) across 10-min post-sonication windows, at 0.625 W/cm^2^ (B; n = 8) and 1.2 W/cm^2^ (C; n = 4). Markers are means ± SD across animals, offset horizontally within each window for legibility; faint points are individual animals. Brackets mark pairs of regions whose relative latencies differ within a window (paired comparisons, Holm-corrected within each window; Table S3).

### 3.7 Forecasting individual gain maps and planning sonication targets

Because the trough follows each animal’s functional map rather than distance from the target, its position on the cortex can be predicted from the map itself. We projected the measured gain profile onto each animal’s pre-sonication PC1 map, allowing local functional gradients to set how far the profile extends in each direction (Sections 2.11 and 2.12). The resulting forecasts follow the irregular geometry of the tonotopic map: the predicted trough stretches along contours of similar response and compresses across steep functional transitions, producing patterns that a radially symmetric model cannot generate (Fig. 5B, D).

**Figure 5.**
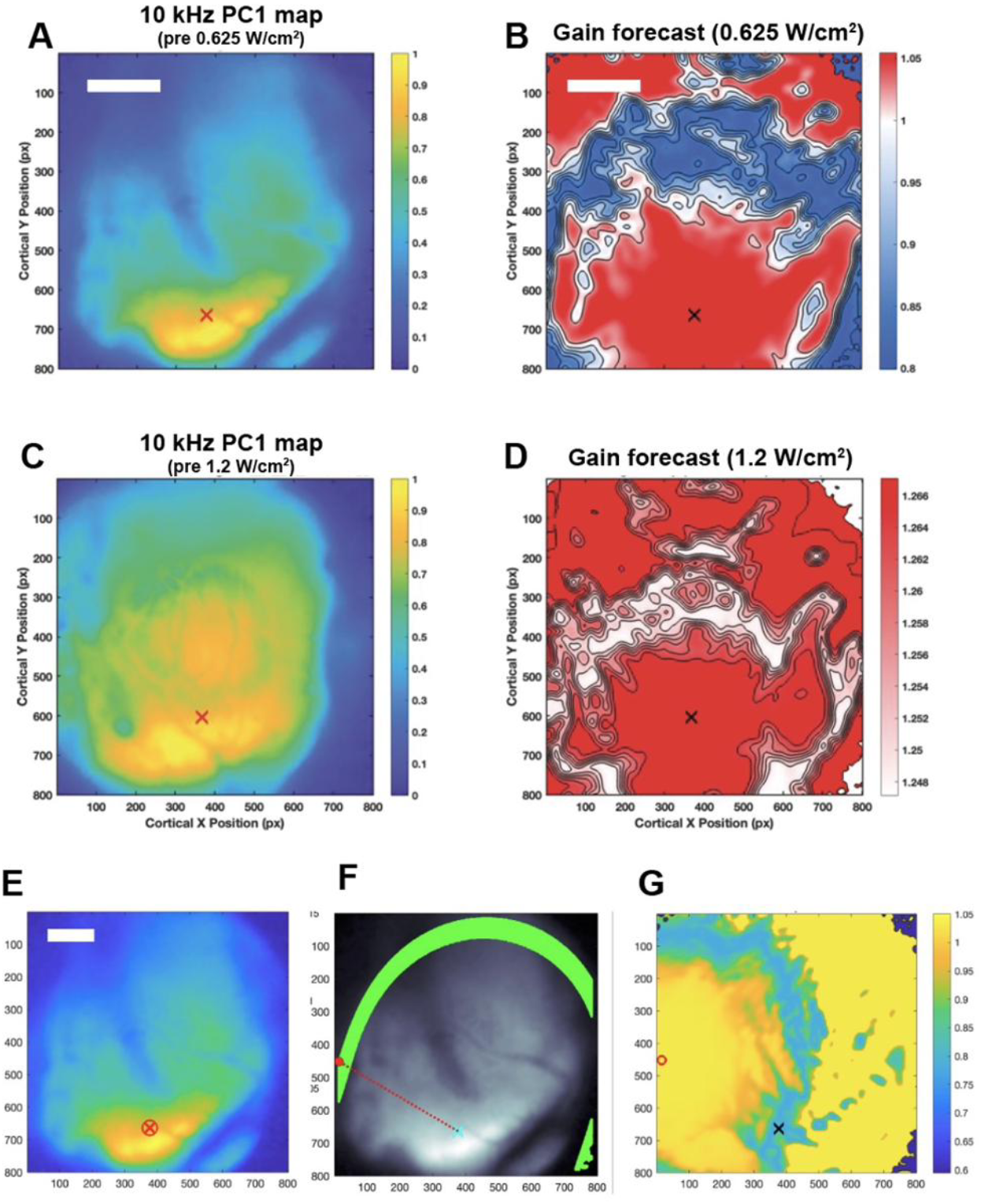
Forecasting an individual animal’s gain map and choosing where to sonicate. (A to D) Gain forecasts for two animals. The pre-sonication 10 kHz PC1 map (A, C; ×, sonication target) determines how the measured spatial gain profile (Fig. S6) is laid out across that animal’s cortex: the profile stretches along contours of similar response and compresses across steep functional transitions, giving the predicted post-sonication gain map at 0.625 W/cm^2^ (B) and 1.2 W/cm^2^ (D). (E to G) Choosing sonication coordinates that produce a specified gain at a chosen cortical location. (E) PC1 map with the chosen location (circled ×). (F) Sonication coordinates predicted to produce the specified gain at that location (green arc); the dotted line links one such coordinate (red) to the chosen location (cyan). (G) Predicted gain map for sonication at the red coordinate in F. Axes are in camera pixels (3.24 μm per pixel). Analysis pipelines are shown in Fig. S4.

The same model, run forward from candidate foci, indicates where to sonicate. For a chosen cortical location and a specified gain at that location, we computed the gain that sonication at each candidate coordinate would deliver there. The coordinates that achieve the specified gain within an error tolerance of ε = max[0.002, 1.15 × min(*E*)] (Section 2.12) form an arc rather than a single point (Fig. 5E, G). Every coordinate on the arc is predicted to produce the same local effect, so the arc offers a choice among targets that differ in their predicted effects elsewhere in the map (Fig. 5F). Planning in the animal’s own functional coordinates thus converts the measured trough into an explicit, testable prescription for where to place the acoustic focus.

### 3.8 Structural connectomic mapping of potential off-target modulation

Given the characteristic elongated axial profile of the FUS beam and the compact dimensions of the mouse auditory cortex, sonication at A1 would be expected to intersect axons entering and leaving the target in addition to mechanoelectrically responsive cells within local cortical tissue. Using the beam-intersection framework described in the Methods (2.14) which leveraged tractography abstractable from anterograde labeling experiments drawn from the Allen Mouse Brain Connectivity Atlas, we determined what share of auditory pathways falls within the sonicated volume, and where the intercepted streamlines terminate (Fig. 6). These included ascending thalamocortical streamlines from the MGB and cortico-cortical streamlines from A1. Because the determination of a true modulatory “threshold” within an FUS beam profile is inherently confounded by its own off-target effects, we performed this assessment over a range of putative intensity thresholds (Table S6).

**Figure 6.**
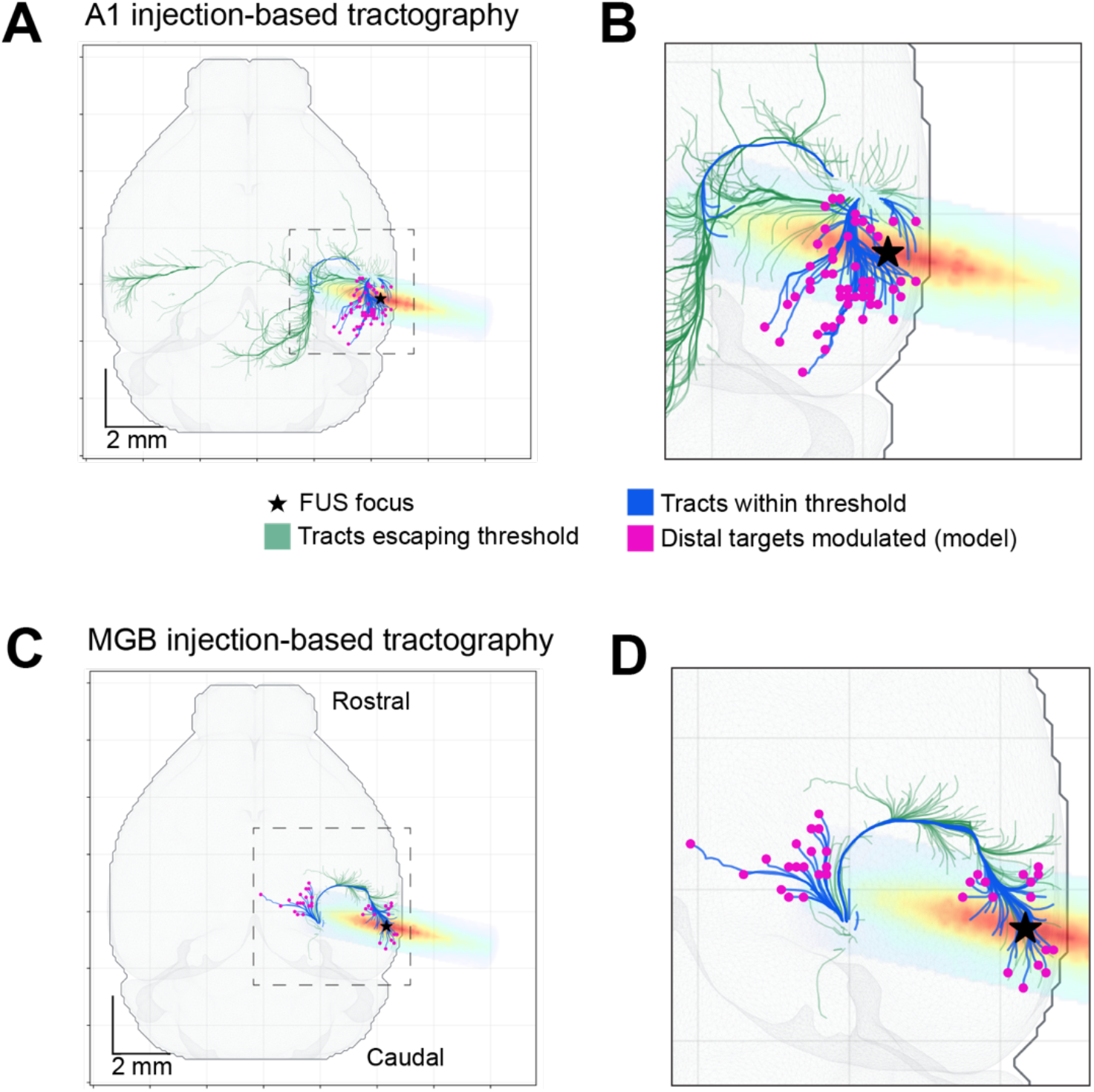
Atlas streamlines intercepted by the 750 kHz beam aimed at right A1. Dorsal views, rostral at top, right hemisphere at right. The jet overlay is the measured beam profile and the star marks the A1 target. (A-B) Allen experiment 120491896 (right A1 injection; 789 streamlines), full view (A) and boxed region (B). (C-D) Allen experiment 180520257 (right MGB injection; 213 streamlines), full view (C) and boxed region (D). Blue: streamlines reaching at least 50% of beam peak (A1 75, MGB 56). Green: streamlines with exposure remaining below 50%. Magenta dots: terminals lying outside the 50% core for streamlines that enter it (A1 53, MGB 35). Streamlines are Allen skeleton components from one experiment per structure, not axons measured in the imaged animals. Scale bars: 2 mm.

Within a volume where local intensity was above 50% of the max pressure, this putative modulatory volume intercepted ∼9% of streamlines whose origins were within A1 and ∼26% of streamlines whose origins were within MGB (Table S4). At this threshold, most intercepted streamlines in both experiments continued past the core to a terminal outside it (A1: ∼7% of all; MGB: ∼16% of all), and the remainder ended within the core of seed regions. Despite anterograde labeling data showing that neurons within A1 send projections terminating on the contralateral hemisphere (green streamlines within Fig. 6), within the hydrophone-measured beam pressure profile for the transducer used in these experiments, all distal terminals outside the core were constrained to the ipsilateral side.

Terminals of streamlines that crossed the core and ended outside it fell in different structures depending on whether tracing was from A1 or MGB (Fig. 6A-B, Table S5). For the MGB anterograde labeling experimental data (Fig. 6C-D, Table S5), distal terminals were in A1, TEa, other auditory cortical areas, thalamus/MGB, none in visual cortex, and structures outside these groups. For the A1 anterograde labeling data, the largest groups were visual cortex and temporal association area, with few in A1 or other auditory cortex, and none in thalamus at the 50% threshold.

## 4. Discussion

### 4.1 Functional topography, not physical distance, organizes FUS-induced gain modulation

In our work, the widefield Ca²⁺ imaging in mouse auditory cortex displayed tonotopic maps that are consistent with prior widefield optical and electrophysiological measurements (Bandyopadhyay et al., 2010; Issa et al., 2014; Romero et al., 2020). Auditory-evoked responses displayed trial-to-trial stochasticity, likely reflecting phase-dependent gating under anesthesia-induced slow-wave activity (Ferron et al., 2009) and possibly rhythmic thalamocortical burst suppression (Kisley and Gerstein, 1999). The functional centroids of individual frequency representations nonetheless remained stable and provided a reliable framework for coordinate-based FUS targeting.

Our central finding is that the spatial footprint of FUS-induced offline gain in mouse A1 is organized by the animal’s own functional cortical topography rather than by physical distance from the sonication focus. The relevant reference frame for describing where an effect occurs is the map itself, defined here by each animal’s pattern of sensory-evoked activity, and not a fixed physical coordinate system referenced to the sonication point. Because functional brackets are oblong and irregular rather than circular (Fig. 2B), a Euclidean analysis pools points that are equidistant from the target but functionally dissimilar together with points that are functionally similar but at different physical distances. That averaging accounts for the Euclidean analysis detecting a change in overall gain without resolving where it occurred.

Spatial specificity in FUS neuromodulation has typically been established by comparing the response at the target with that at one or more displaced sites, or by relating the response to the modeled acoustic field. This approach has demonstrated focal effects at centimeter scale with EEG in humans (Mueller et al., 2014), in mice with c-Fos expression (Kim et al., 2021), and at sub-millimeter scale with two-photon Ca²⁺ imaging in awake mouse visual cortex (Cheng et al., 2022). Such comparisons show that an effect is confined to roughly the dimensions of the focus, but they do not reveal how the magnitude or sign of the effect varies across the tissue within and around it. Studies that have mapped FUS effects across whole regions find that the affected territory does not match the acoustic field. Offline sonication of the macaque amygdala alters functional coupling at the target and in anatomically connected regions that lie outside the beam (Folloni et al., 2019). In mouse, sonication-induced vasoconstriction spares hippocampus but extends across thalamus, hypothalamus and cortex, and does not scale with the local change in neural activity (Murphy et al., 2024). In mouse somatosensory cortex, we previously found that FUS alters the spatial pattern, and not necessarily just the amplitude, of a sensory-evoked response at the focus (Fisher and Gumenchuk, 2018). The present study measures gain continuously across a cortical map and expresses it in the subject’s own functional coordinates, the frame in which the structure reported here appears.

The geometry we observe is consistent with the intrinsic circuitry of A1. Intracortical connectivity in mouse auditory cortex is patchy and oriented with respect to the tonotopic axis rather than distributed isotropically: supragranular intra-laminar connections are elongated along the isofrequency axis, local circuits are asymmetric with respect to that axis, and functionally connected patches are organized by the spectral relationship between sites rather than by their separation (Jendrichovsky et al., 2025; Oviedo et al., 2010; Watkins et al., 2014). The same organizing principle holds in species with substantially larger auditory cortices; in the cat, patchy intrinsic connections link iso-tuned subregions (Read et al., 2001), and in the guinea pig, tone-evoked activity propagates preferentially along isofrequency contours rather than radially in guinea pig (Song et al., 2006). Absolute distances, however, are not transferable between these preparations. The tonotopic gradient occupies 1 to 2 mm in mouse (Tsukano et al., 2016) and a substantially greater extent in cat and guinea pig, so a connectional span measured in those species covers a different fraction of the map than the same figure would in mouse, and should not be read as matching the locus of the modulatory kernel we observed in the present work. Prior work therefore supports the orientation and shape of the modulatory footprint rather than its radius: connectivity laid out with respect to the map provides a basis for an effect that stretches along contours of similar response and compresses across steep functional transitions. This is the geometric principal that our forward model reproduces (Section 3.7).

### 4.2 Mechanistic constraints from the time course and spatial profile

Two of this study’s key experimental findings—the 10–20 min emergence of a persistent modulatory trough and the annular, FUS intensity-dependent spatial footprint of gain—bear significant alignment with prior work. The bimodal sign of the effect at different intensities, for example, may be accounted for by differential recruitment of inhibitory and excitatory elements. In terms of acute neuronal effects, low-intensity FUS has been shown to engage parvalbumin-positive interneurons in mouse cortex (Kim et al., 2026; Murphy et al., 2022), fast-spiking and regular-spiking neurons respond differently to acoustic parameters (Yu et al., 2021), and parameter sets that excite and inhibit are separable across cell types and targets (Murphy et al., 2024). However, because these acute electrophysiological effects relax within minutes, it is unlikely they alone are responsible for preserving gain modulation 40 min later. More likely, neuronal substrates for acute perturbations transition to persistent time courses through synaptic plasticity and intrinsic excitability changes. For example, FUS induces hippocampal long-term depression (LTD), causing field EPSP (fEPSP) slope reductions that persist over 20–60 min (Niu et al., 2022). Complementing synaptic LTD, FUS drives long-lasting increases in intrinsic excitability via ionotropic glutamate receptor-dependent dephosphorylation of voltage-gated potassium channel subunits *K_v_*2.1 (Clennell et al., 2023, 2021), alongside sustained reductions in cortical GABA levels in humans (Yaakub et al., 2023).

A threshold in a slowly expressed effector could account for the radius without invoking a separate spatial mechanism. Because intensity falls off smoothly from the focus, a dose positioned near a regime boundary at the focus necessarily falls on the other side of that boundary at lager radii. Beyond neuronal considerations, astrocytes are recruited by ultrasound at lower acoustic pressures than neurons, mounting strong Ca²⁺ responses that drive sustained, cyclooxygenase-dependent vascular changes over a territory exceeding the focus (Murphy et al., 2024). This astrocytic sensitivity operates alongside broader non-neuronal cascades: FUS elevates intracellular Ca²⁺ in neurons, astrocytes, endothelial cells, and pericytes through mechanosensitive and transient receptor potential channels rather than store release (Newman et al., 2023). As a common upstream signal, Ca²⁺ engages cyclooxygenase, nitric oxide, and transcriptional pathways capable of converting brief acoustic exposure into lasting functional states. Although Murphy et al. (2024) observed vasoconstriction, our earlier measurements with distinct parameters revealed persistent vasodilation (Shen et al., 2025), indicating that vascular sign may be protocol-dependent. Because cortical hemodynamics ordinarily follow neural activity rather than drive it (Iadecola, 2017), these microvascular responses are best interpreted as functional reporters of shared astrocytic and vascular effectors rather than primary drivers of neural gain. Finally, non-reactive neuroimmune signaling offers a complementary pathway, as glial tumor necrosis factor α regulates synaptic strength via AMPA-receptor trafficking (Stellwagen and Malenka, 2006) and astrocytes instruct microglial state (McAlpine et al., 2021), processes representing physiological modulation rather than reactive gliosis, consistent with the absence of barrier disruption or tissue damage at 1.2 W/cm² (Fig. S3).

Two additional, non mutually exclusive routes may bear on the footprint. First, because the focal volume extends axially beneath the targeted tonotopic locations on A1, our exploratory streamline intersection (Fig. 6, Table S4) indicates that in addition to cortico-cortical fibers leaving A1, ascending thalamocortical fibers from the MGB are intercepted; these MGB pathways terminate across A1, adjacent auditory fields, temporal association area (TEa), and thalamus. By contrast, intercepted A1 streamlines passing through the focal core terminate outside A1, in TEa, and visual cortex. This suggests that long-range cortico-cortical overlap may not by itself account for the intra-A1 trough in gain and that modulation of thalamocortical fibers in passage could plausibly contribute to the broader amplification seen at 1.2 W/cm² and to remote, connectivity-patterned effects outside the imaging field (Folloni et al., 2019). Separately, the shift in relative latency between the PC1, PC2 and PC3 regions speaks to propagation rather than amplitude, since these fields receive parallel inputs from distinct subdivisions of the medial geniculate body and differ systematically in onset latency (Hackett et al., 2011; Tsukano et al., 2017, 2016; Wang et al., 2024). Perturbing a subset of these ascending MGB streamlines in passage offers a plausible structural route for shifting relative timing across fields. Discriminating among these accounts could be addressed through cell-type-specific Ca²⁺ reporting (Murphy et al., 2022), two-photon imaging during sonication to locate the intensity at which the sign of the effect reverses (Cheng et al., 2022), and potentially whole-brain imaging of c-Fos expression, e.g., using light-sheet microscopy (Kirst et al., 2020), which can report recent brain-wide changes in neuronal activity.

### 4.3 Translational implications for chronic tinnitus and sonication planning

Coordinate-based, non-invasive sensory map editing has direct relevance to chronic conditions in which map organization itself is pathological. Chronic tinnitus is a leading example: it has been the most prevalent service-connected disability among US military veterans since 2007, with 2.9 million veterans compensated for it in fiscal year 2023 (US Department of Veterans Affairs, 2023). Tinnitus arises from a number of conditions, including age-related hearing loss (Elgoyhen et al., 2015), but one of its most common manifestations, and the predominant service-related one, is a maladaptive plastic remapping in which peripheral hearing loss leads to aberrant hyperexcitability and over-representation of edge frequencies in A1 (Eggermont and Roberts, 2004).

Existing non-invasive interventions, including transcranial direct current stimulation and bimodal sound-plus-stimulation approaches, have produced limited and inconsistent lasting benefit, potentially in part because they cannot address discrete, frequency-specific cortical territory (Conlon et al., 2020; Jeon et al., 2023; Langguth, 2020). Our results show that FUS can place and constrain an offline gain change at sub-millimeter scale within the compressed mouse auditory cortex, 1 to 2 mm² in total. The human tonotopic core in Heschl’s gyrus spans a substantially larger gradient of approximately 27 to 30 mm across seven octaves. If a focal acoustic profile on the order of 0.5 to 1.0 mm can be maintained through the human skull, a clinician could in principle identify the pathologically shifted coordinates of a patient’s own frequency map and deliver a tailored sonication protocol to that location, informed by a personalized functional map in the same way that sonication was targeted to each mouse’s own PC1 centroid here.

Applying this approach in patients would require control of the acoustic field beyond the single-focus sonication used here. Holographic transcranial FUS forms customized pressure patterns without scanning a focus point by point, and has been demonstrated transcranially in rodents with an acoustic design extended to the human skull (Estrada et al., 2026). In that work, foci placed 0.5 to 1 mm apart recruited shared circuitry, producing a combined response larger than the sum of the individual foci and lowering the activation threshold below that of a single focus of equal total power. Because multi-focal effects are not additive, extending the planning approach of Section 3.7 to multi-focal patterns would require measuring the gain profile for those patterns directly rather than summing single-focus profiles. In parallel, utilizing patient-specific diffusion tensor imaging (DTI), lightweight beam-intersection screening of the kind outlined here could complement functional planning by helping compare candidate transducer geometries and anticipate off-target tract exposure prior to intervention. Any such protocol must also be tied to perception. Whether the macroscopic re-weighting of tonal representations reported here corresponds to a perceptible shift in hearing thresholds can be tested with established head-fixed paradigms for tone detection and two-tone frequency discrimination in mice, in which cortical manipulation has already been shown to bias auditory choice (Ceballo et al., 2019; Francis et al., 2018).

### 4.4 Limitations and future directions

Limitations of the current study include both systematic effects and generalizability concerns. As described in Section 3.8, the axial extent of the beam delivered non-negligible acoustic exposure to thalamic nuclei and ascending auditory fibers, so a direct thalamic or thalamocortical contribution cannot be ruled out. Transducer geometries that constrain the focus axially, such as crossed-beam configurations that achieve an approximately 1 mm focus in all three dimensions (Kim et al., 2021), could remove this confound; in animal experiments, higher fundamental frequencies offer an alternative route to a smaller focal volume (Cheng et al., 2022). A second limitation is that the framework is at present fit to and validated against the same measurements from which it was derived. Prospective validation against new, held-out sonication targets, ideally combined with an independent behavioral readout, is needed before it can be used to plan interventions.

A third limitation concerns the mechanistic accounts of Sections 4.3 and 4.4. The measurements reported here constrain those accounts by their time course and spatial profile but do not discriminate among them, since no cell-type-specific, ionic, vascular or immune readout was obtained in the same preparation and over the same window as the gain measurements. The reporter used labels predominantly excitatory neurons, so the activity of interneurons, astrocytes and microglia was not observed. The experiments proposed in Sections 4.3 and 4.4, cell-type-specific photometry, recording of the slow cortical potential and extracellular K⁺ at the focus and at the trough radius, and assessment of microglial state at the relevant radii and timepoints, are each designed to separate these candidates and are the logical next step.

## CRediT authorship contribution statement

**Jacob R. Hehir**: Writing, review and editing; Writing, original draft; Visualization; Investigation; Formal analysis; Data curation; Conceptualization. **YuBing Yolanda Shen**: Writing, review and editing; Validation; Investigation; Formal analysis. **Claire J. Lee**: Writing, review and editing; Formal analysis, Validation. **Jonathan A. N. Fisher**: Writing, review and editing; Writing, original draft; Supervision; Resources; Project administration; Funding acquisition; Formal analysis; Conceptualization.

## Funding

This work was supported by award R01EB028319 from the National Institute of Biomedical Imaging and Bioengineering to J.A.N.F.

## Declaration of competing interest

The authors have no conflict of interest related to this work.

## Acknowledgments

The authors thank Furong Hu for assistance with immunohistological analysis, tissue preparation and related imaging; Aparna Singh and Elisa Konofagou for assistance calibrating ultrasound systems; and CheukHimIan Lam for assistance with image processing. Grammar and phrasing were edited with the assistance of Microsoft Copilot and Claude (Anthropic). No AI tools were used for the generation or interpretation of results, nor in experimental design.

## Data availability

The datasets generated and analyzed in this study are available from the corresponding author upon reasonable request.

## Code availability

Custom code used for analysis in this study, including the connectomics and beam-intersection analysis, is available on GitHub and access will be provided upon request to the corresponding author.

## Supplementary Materials

**Figure S1.**
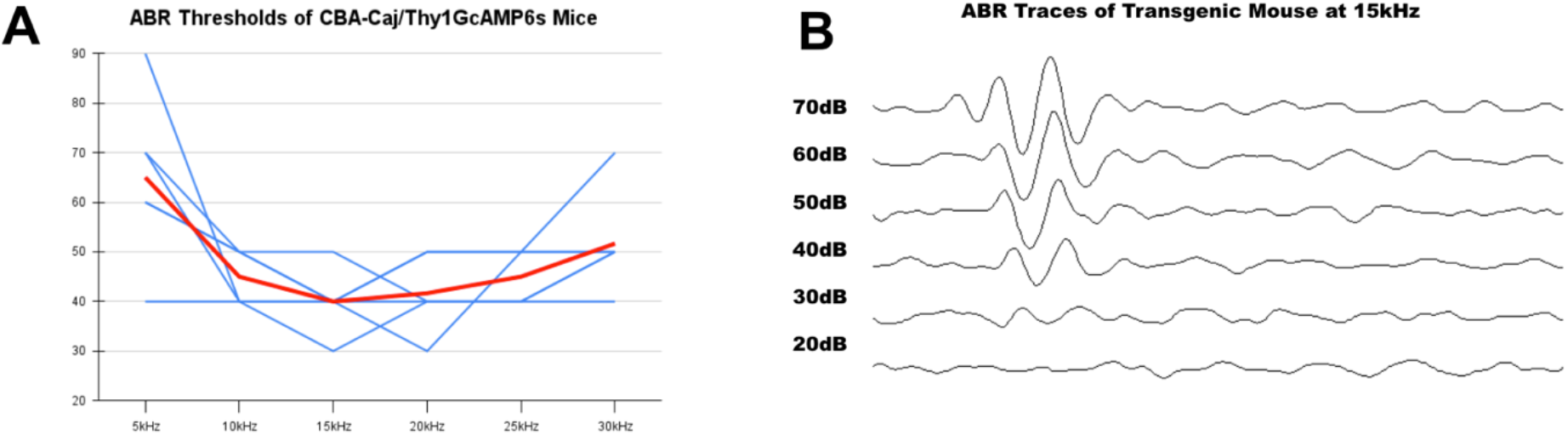
Hearing thresholds in Thy1-GCaMP6s × CBA/CaJ mice. (A) Auditory brainstem response (ABR) thresholds from 5 to 30 kHz for individual animals (blue) and their mean (red). (B) ABR waveforms evoked by a 15 kHz tone pip from 70 to 20 dB SPL in one animal; the threshold for this animal at 15 kHz is 30 dB SPL.

**Figure S2.**
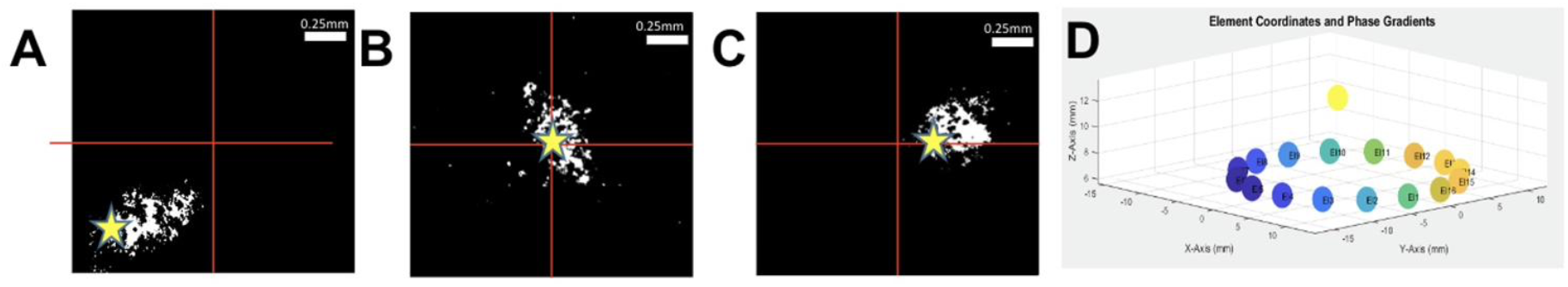
Beam steering and targeting validation. (A to C) Thermochromic film response to FUS steered to three target coordinates (yellow stars): x = −0.5, y = −0.5 mm (A); x = 0, y = 0 mm (B); x = 0.25, y = 0 mm (C). Heated regions appear white; red crosshairs mark the center of the imaging field. Scale bars, 0.25 mm. (D) Element positions and per-element phase gradients used to steer the 16-element array to the focus (yellow dot); the beam propagates along +z.

**Figure S3.**
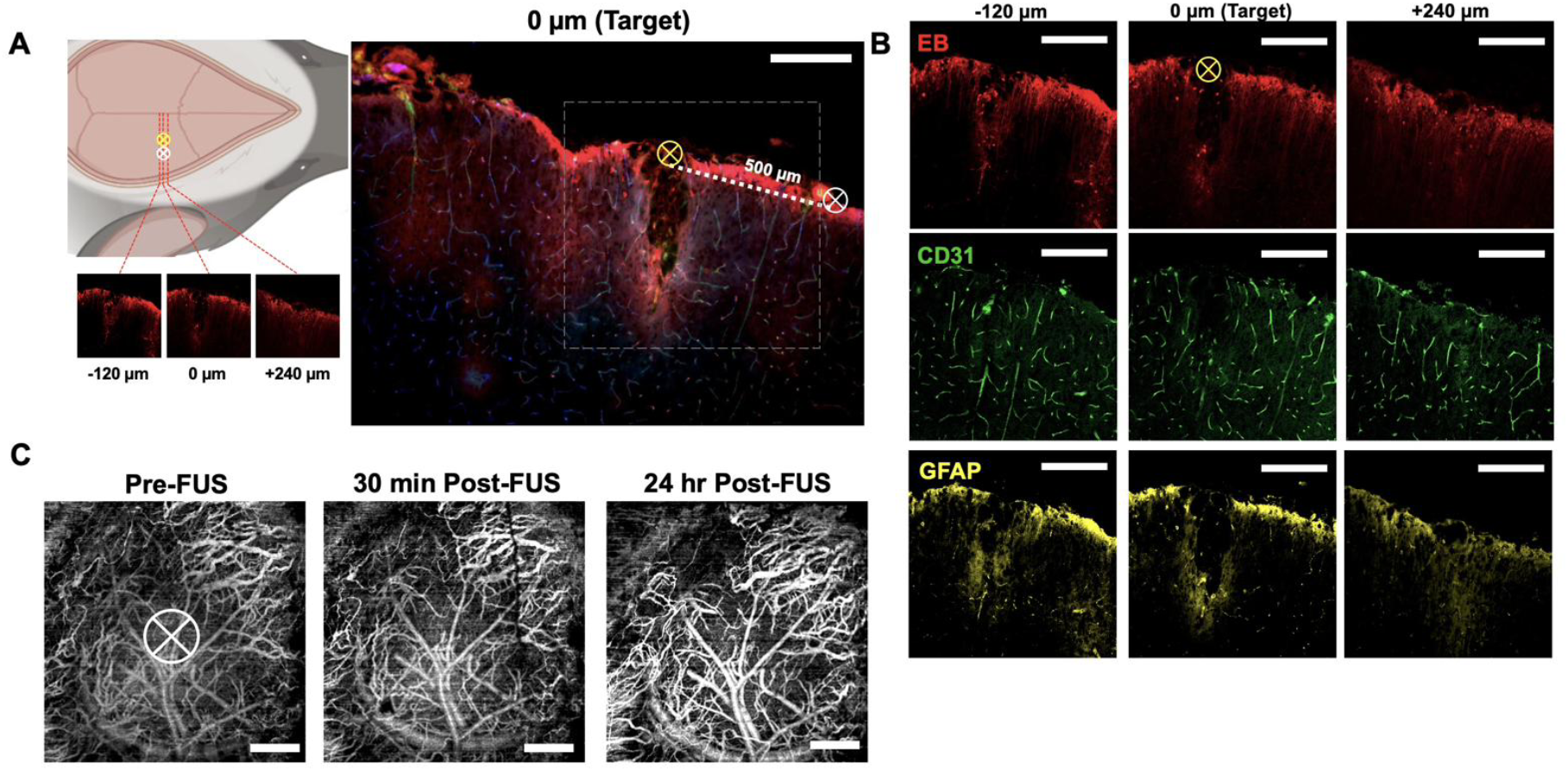
Histological and angiographic safety assessment. (A) Left: sonication sites. A high-intensity marker dose (223 W/cm^2^ *I*_sppa_, 100% duty cycle; yellow crosshair) was delivered 0.5 mm from a site sonicated at 1.2 W/cm^2^ (50% duty cycle; white crosshair), and sections were taken at the target and at −120 μm and +240 μm along the rostrocaudal axis. Right: section through both sites, showing a lesion and Evans blue extravasation (red) at the marker dose and no tissue change at the 1.2 W/cm^2^ site; the dashed line spans the 500 μm between them. (B) Evans blue (blood-brain barrier integrity), CD31 (vascular endothelium) and GFAP (astrocytes) at the three section positions. (C) OCT angiography before, 30 min after and 24 h after sonication at 1.2 W/cm^2^ (target circled), showing no change in the microvasculature. Scale bars, 250 μm (A, B) and 500 μm (C).

**Figure S4:**
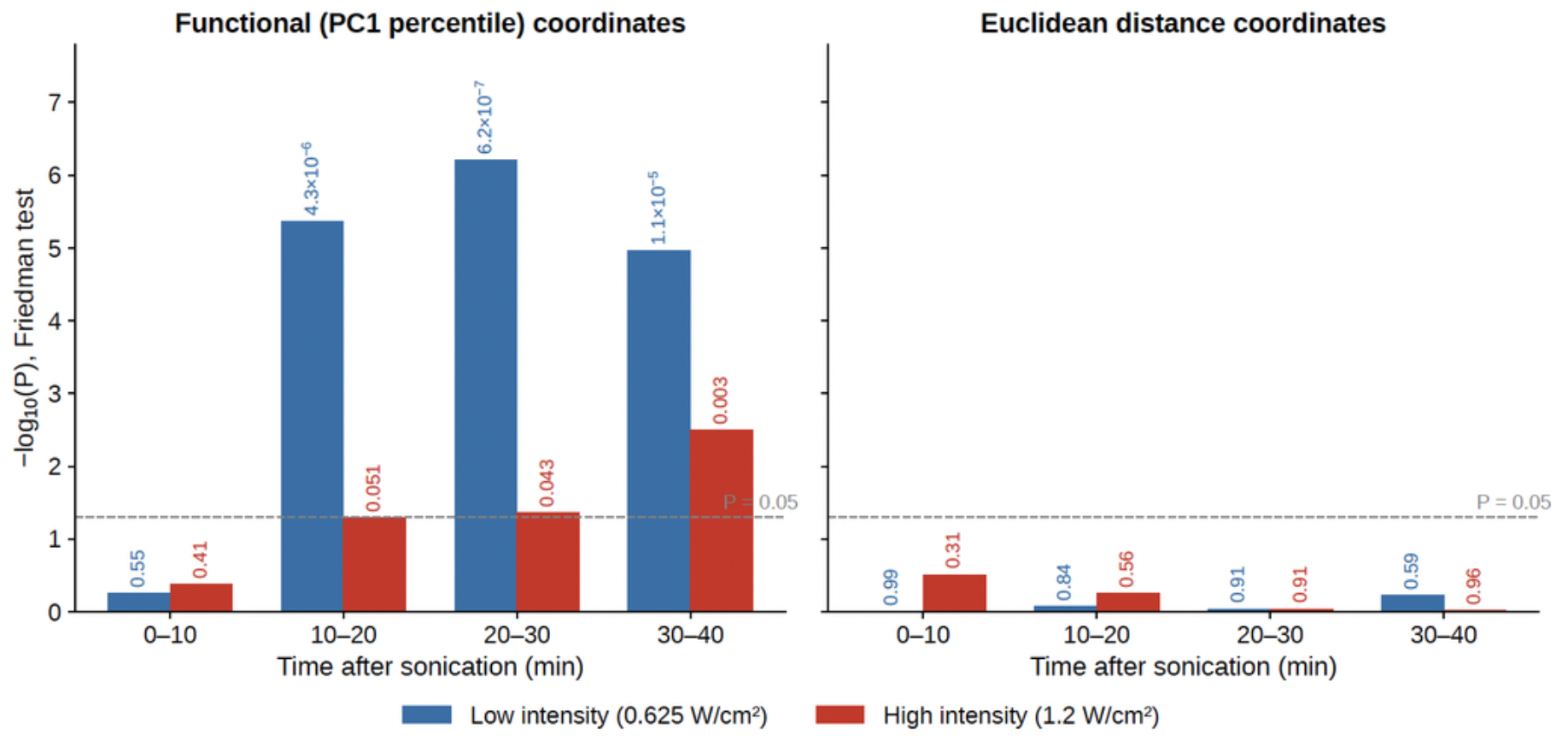
Omnibus test for spatial structure. For each post-sonication window and intensity, a Friedman repeated-measures test of the null hypothesis that gain does not differ across locations within animals, with animals as blocks and location as the within-animal factor. Locations were defined either by percentile of the pre-sonication PC1 map (left; 9 brackets, df = 8) or by Euclidean distance from the sonication target (right; 5 rings of 250 μm, df = 4). Bars show −log_10_(P), with the exact P above each bar; the dashed line marks P = 0.05. One test was performed per window, intensity and coordinate system, and no location was selected before testing.

**Figure S5.**
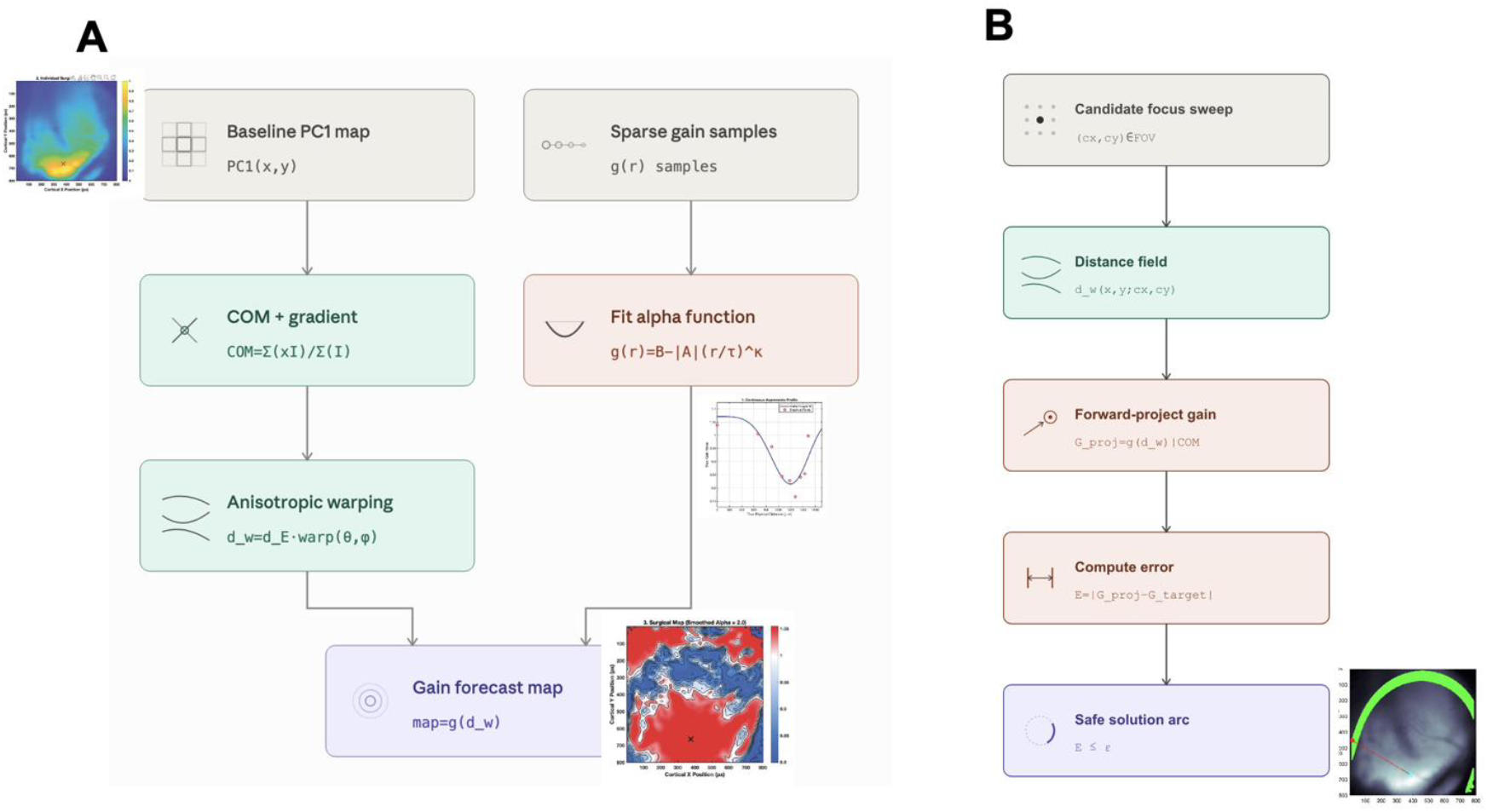
Analysis pipelines for the gain forecasts and sonication planning in Fig. 4. (A) Forecast: the center of mass and local gradient of the pre-sonication PC1 map define an anisotropic distance field (Section 2.11), along which the spatial gain profile fitted to the measured bracket gains is evaluated to give a predicted gain map. (B) Planning: for each candidate sonication coordinate, the predicted gain at the chosen location is compared with the specified gain, and coordinates within tolerance form the solution arc (Section 2.12).

**Figure S6.**
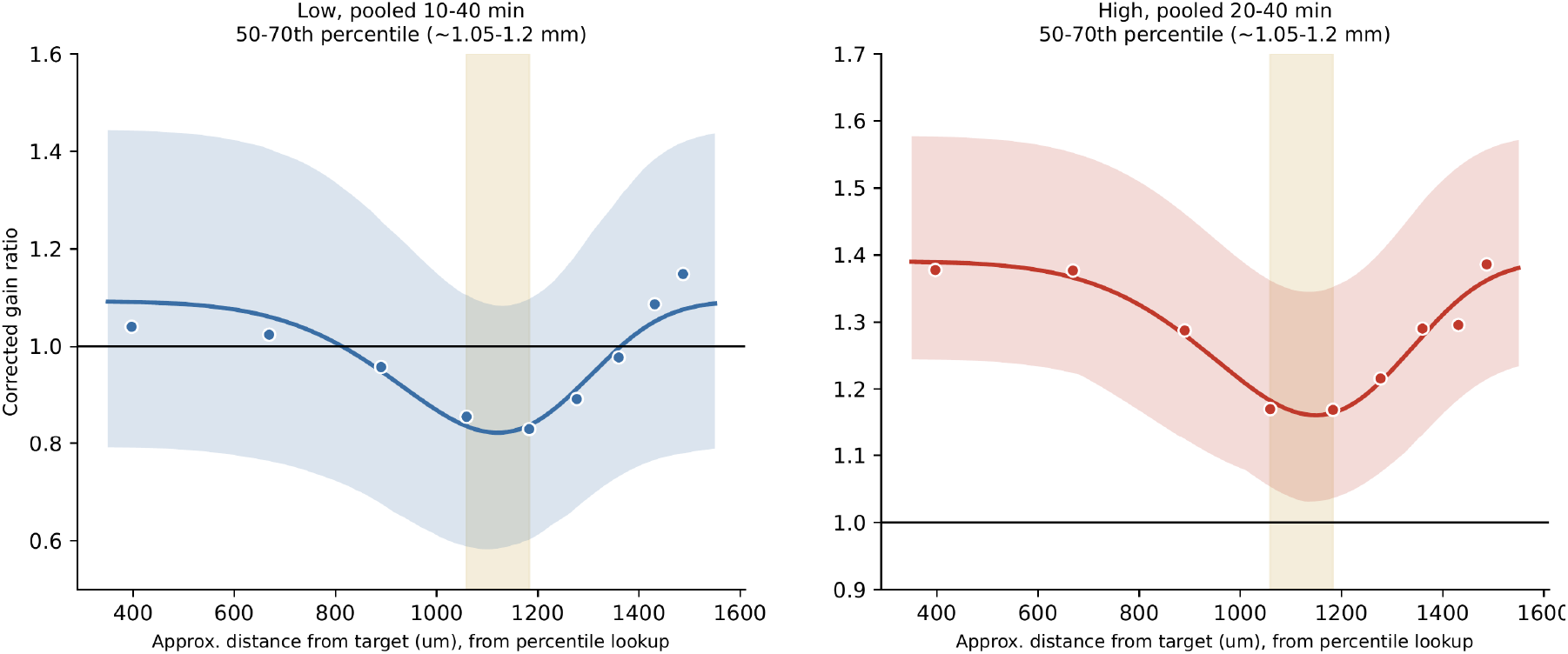
Spatial profile of offline gain. Gain in each functional-percentile bracket (points, group means) plotted against the bracket’s mean distance from the sonication target (Fig. 2C), with the fitted profile (line; Section 2.11) and its 95% confidence band (shading), at 0.625 W/cm^2^ pooled over 10 to 40 min (left) and 1.2 W/cm^2^ pooled over 20 to 40 min (right). Pooled windows are those in which the Friedman test detected spatial structure (Fig. 2F). The vertical band marks the 50th to 70th percentile brackets (about 1.05 to 1.2 mm). Fit parameters are given in Table S2.

**Table S1.** Mean distance from the PC1 center of mass for each functional-percentile bracket, by group.

| **PC1 bracket** | **Control (n=3)** | **Low FUS, 0.625 W/cm<sup>2</sup>** | **High FUS, 1.2 W/cm<sup>2</sup>** |
|---|---|---|---|
| 80 to 90% | 637 μm | 573 μm | 606 μm |
| 70 to 80% | 922 μm | 796 μm | 913 μm |
| 60 to 70% | 1,130 μm | 1,045 μm | 1,131 μm |
| 50 to 60% | 1,278 μm | 1,225 μm | 1,307 μm |

**Table S2.** Location and depth of the fitted spatial gain profile, by intensity and post-sonication window. τ is the distance from the sonication target at which the fitted profile reaches its minimum. Distances are converted from functional percentile using the group-mean bracket distances. Pooled fits use the 10-min windows in which the Friedman test detected spatial structure (Fig. S4). Relative trough depth is expressed against each fit’s core gain *B*; trough gain is reported for pooled fits only. 0.625 W/cm^2^, n = 8; 1.2 W/cm^2^, n = 4.

| **Fit** | **n** | **τ, mm (95% CI)** | **Core gain *B*** | **Trough gain (relative depth)** |
|---|---|---|---|---|
| 0.625 W/cm<sup>2</sup>, 10 to 20 min | 8 | 1.06 (0.76 to 1.14) | 1.06 |  |
| 0.625 W/cm<sup>2</sup>, 20 to 30 min | 8 | 1.16 (1.04 to 1.20) | 1.13 |  |
| 0.625 W/cm<sup>2</sup>, 30 to 40 min | 8 | 1.08 (0.69 to 1.17) | 1.16 |  |
| 0.625 W/cm<sup>2</sup>, pooled 10 to 40 min | 8 | 1.12 (1.10 to 1.14) | 1.09 | 0.82 (−25%) |
| 1.2 W/cm<sup>2</sup>, 20 to 30 min | 4 | 1.18 (1.15 to 1.18) | 1.30 |  |
| 1.2 W/cm<sup>2</sup>, 30 to 40 min | 4 | 1.12 (1.05 to 1.16) | 1.50 |  |
| 1.2 W/cm<sup>2</sup>, pooled 20 to 40 min | 4 | 1.15 (1.10 to 1.17) | 1.39 | 1.16 (−17%) |

**Table S3.** Paired comparisons of peak latency between PC regions, by intensity and post-sonication window. Cells give raw P / Holm-corrected P (m = 3 within each window and intensity). Low intensity n = 8, except n=6 for PC1 vs PC2 and PC2 vs PC3 in the 30 to 40 min window; high intensity n = 4. None of the 24 comparisons of a region against its own pre-FUS baseline reached significance at either intensity, with or without correction (smallest raw P=0.075).

| **Comparison** | **0 to 10 min** | **10 to 20 min** | **20 to 30 min** | **30 to 40 min** |
|---|---|---|---|---|
| Low, PC1 vs PC2 | 0.014 / 0.041 | 0.088 / 0.211 | 0.213 / 0.213 | 0.353 / 1.000 |
| Low, PC2 vs PC3 | 0.032 / 0.064 | 0.070 / 0.211 | 0.017 / 0.050 | 0.771 / 1.000 |
| Low, PC1 vs PC3 | 0.909 / 0.909 | 0.382 / 0.382 | 0.093 / 0.186 | 0.765 / 1.000 |
| High, PC1 vs PC2 | 0.423 / 0.967 | 0.188 / 0.565 | 0.423 / 1.000 | 0.423 / 1.000 |
| High, PC2 vs PC3 | 0.423 / 0.967 | 0.423 / 0.845 | 0.423 / 1.000 | 0.568 / 1.000 |
| High, PC1 vs PC3 | 0.322 / 0.967 | 0.423 / 0.845 | 0.903 / 1.000 | 0.705 / 1.000 |

**Table S4:** Streamlines intercepted at 50% of peak (percent of all streamlines in each experiment).

|  | **A1 (120491896)** | **MGB (180520257)** |
|---|---|---|
| Total streamlines | 789 | 213 |
| Enter 50% core | 75 (9.5%) | 56 (26.3%) |
| Enter core, terminal outside | 53 (6.7%) | 35 (16.4%) |
| Enter core, terminal inside | 22 (2.8%) | 21 (9.9%) |
| Contralateral terminals (outside core) | 0 | 0 |

**Table S5.**
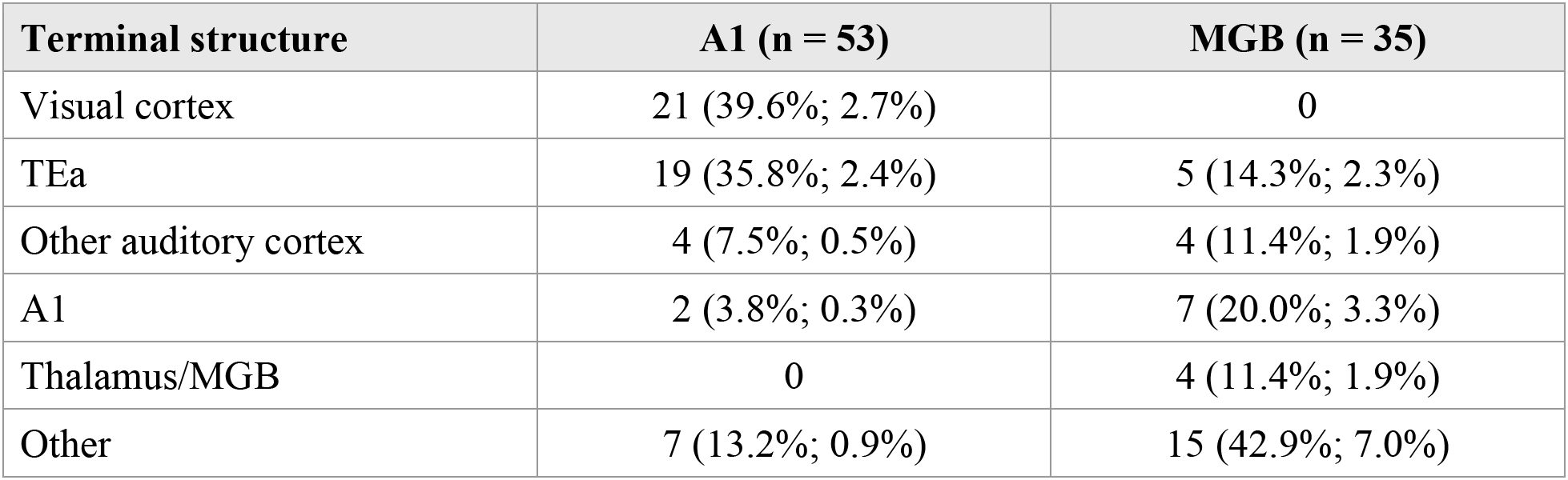
Terminal structure of streamlines that enter the 50% core and end outside it. Percentages describe destinations of this subset only, not of all streamlines. TEa = temporal association area; MGB = medial geniculate body.

| **Terminal structure** | **A1 (n = 53)** | **MGB (n = 35)** |
|---|---|---|
| Visual cortex | 21 (39.6%; 2.7%) | 0 |
| TEa | 19 (35.8%; 2.4%) | 5 (14.3%; 2.3%) |
| Other auditory cortex | 4 (7.5%; 0.5%) | 4 (11.4%; 1.9%) |
| A1 | 2 (3.8%; 0.3%) | 7 (20.0%; 3.3%) |
| Thalamus/MGB | 0 | 4 (11.4%; 1.9%) |
| Other | 7 (13.2%; 0.9%) | 15 (42.9%; 7.0%) |

**Table S6.** Intercepted streamlines as a function of beam contour.

| **Threshold (% of peak)** | **A1** | **MGB** |
|---|---|---|
| 20 | 620 / 789 (78.6%) | 213 / 213 (100%) |
| 30 | 616 / 789 (78.1%) | 211 / 213 (99.1%) |
| 40 | 612 / 789 (77.6%) | 200 / 213 (93.9%) |
| 50 | 75 / 789 (9.5%) | 56 / 213 (26.3%) |
| 60 | 53 / 789 (6.7%) | 19 / 213 (8.9%) |
| 70 | 45 / 789 (5.7%) | 13 / 213 (6.1%) |
| 80 | 28 / 789 (3.5%) | 5 / 213 (2.3%) |
| 90 | 14 / 789 (1.8%) | 1 / 213 (0.5%) |
| 100 | 0 | 1 / 213 (0.5%) |

## Notes

### Competing Interest Statement

The authors have declared no competing interest.

